# Cholesteryl esters and high protein-to-lipid ratios distinguish Non-Vesicular Extracellular Particles from Extracellular Vesicles

**DOI:** 10.1101/2025.07.01.662358

**Authors:** Sayam Ghosal, Rita Leporati, Árpád Varga, Petra Susanszki, Nóra Fekete, Tamás László, Tünde Bárkai, Fülöp Károly Grébecz, Delaram Khamari, Tárek Zoltán Magyar, Marcus Hoering, Krisztina V Vukman, Bernadett R Bodnár, Brachyahu M. Kestecher, Mohamed A Fattah, Csaba Bödör, József Maléth, Gerhard Liebisch, Evelyn Orsó, Edit I Buzas, Xabier Osteikoetxea

## Abstract

Extracellular vesicles (EVs) are central to intercellular communication, yet the mechanisms underlying their biogenesis and diversity remain incompletely understood. Here, we integrate meta-analysis, advanced lipidomic, protein-to-lipid profiling, and super-resolution imaging to define the fundamental principles governing EV heterogeneity. Our meta-analysis of published transmission electron microcrographs across kingdoms reveals a highly conserved 110 nm average diameter and 200 nm upper size limit for intraluminal vesicles (ILVs), which are secreted as exosomes. Besides classical EV populations, we also characterize a distinct nanoparticle class: 167000 xg pellet of non-vesicular extracellular particles (167k-NVEPs), which exhibit a significantly higher protein-to-lipid ratio than 14000 xg pellet of large EVs (14k-lEVs) and 100000 xg pellet of small EVs (100k-sEVs), as measured by both biochemical assays and Raman spectroscopy. Lipid profiling demonstrates that 167k-NVEPs exhibit significant enrichment in cholesteryl esters and triacylglycerols, lipids typically associated with lipid droplets and the endosome/lysosome system. Analysis of lipid carbon-chain lengths reveals distinct signatures: 167k-NVEPs show pronounced enrichment at 16 and 18 carbons, while 100k-sEVs display enrichment at 32 and 34 carbons. This divergence indicates a potential connection to flexible biogenesis pathways. Marker heterogeneity across EV populations, confirmed by confocal and super-resolution microscopy, further underscores the limitations of relying on canonical tetraspanins for EV classification. Notably, 167k-NVEPs (likely exomeres) exhibit enrichment of Arf6 and CD63. Together, our findings provide compositional, biophysical, and molecular evidence supporting the formal recognition of 167k-NVEPs as a distinct class of extracellular particles and enabling exploring in disease biology and therapeutic delivery.

**Significance Statement:** Extracellular vesicles (EVs) are critical mediators of intercellular communication, yet their classification remains clouded by ambiguity in terms of their composition and biogenesis. This study resolves critical uncertainties through a cross-kingdom meta-analysis, establishing a conserved ∼110nm diameter and ∼200 nm upper size limit for intraluminal vesicles (ILVs), the precursors to exosomes. More significantly, we identify non-vesicular extracellular particles (167k-NVEPs) as a distinct class based on their unique sterol-rich lipidome, enrichment in lipids of 16 and 18 carbon chain length, elevated protein-to-lipid ratio, and functional cargo delivery. These features, alongside evidence of non-canonical origin and functional cargo delivery, establish NVEPs as a discrete class of extracellular particles.

## Introduction

The field of extracellular vesicle (EV) research has progressed significantly in recent decades, elucidating the complex nature and increasing potential of EVs for disease diagnosis and therapy (1). EVs have demonstrated dual functions, influencing both the progression and inhibition of various conditions including cancer (2), cardiovascular disorders (3, 4), and neurological diseases (5). Current understanding categorizes EV biogenesis broadly into two primary pathways: direct budding from the plasma membrane (ectocytosis) or the formation of intraluminal vesicles (ILVs) within multivesicular bodies (MVBs), which are subsequently released into the extracellular space as exosomes (exocytosis) (6). This diverse origin results in a heterogeneous population of EVs, occasionally with relatively defined small or large sizes, leading to substantial debate regarding their precise sources. Small EVs (∼50-200 nm diameter, sEVs) are hypothesized to be primarily secreted through the exosomal pathway, while large EVs (>200 nm diameter, lEVs) have been postulated to be predominantly released by ectocytosis (7). Further complexity in biogenesis was recently introduced with the discovery of non-vesicular extracellular particles (NVEPs) such as exomeres (8) and supremeres (9), whose biogenesis mechanisms remain unknown. NVEPs merit broad scientific focus because they are highly abundant, carry disease-linked proteins and RNAs once ascribed to EVs, and play key roles in intercellular signaling and clinical applications (10, 11). Accurately profiling NVEP is essential to decipher cellular secretory pathways and to tailor biomarker discovery and therapeutic delivery, marking this effort as a pivotal frontier in cell biology and translational medicine.

Questions persist as to whether NVEPs follow conventional EV release pathways or arise through distinct biogenetic mechanisms. Their molecular composition and origin remain poorly defined, largely due to the absence of systematic, multiparametric comparisons with canonical EV subtypes. Given these unresolved issues, continued investigation into the origin, structure, and function of both EVs and NVEPs is essential to refine current classification frameworks and fully understand their biological roles. In this study, we address these questions by comprehensively characterizing distinct EV subpopulations, including lEVs, sEVs, and NVEPs. Based on differential centrifugation and filtration, we sequentially isolated two EV pellets; first a pellet at 14,000 x g enriched in lEVs (14k-lEVs) and a second pellet at 100,000 x g enriched in sEVs (100k-sEVs), followed by a third pellet collected at 167,000 x g enriched in NVEPs (167k-NVEPs) as has been shown earlier (9, 12–14). Aiming to resolve the debate about EV size and its correlation with their biogenesis, we performed a cross-kingdom comprehensive meta-analysis to observe whether EVs with a diameter greater than 200 nm can origin from the ILV route. Orthogonal, multiparametric characterization was also performed to compare NVEPs alongside canonical EV subtypes, assessing their morphology, protein to lipid ratio, single particle characterization, Raman spectroscopy, lipidomic analysis, high-resolution flowcytometry and super resolution microscopy. To trace the origin of these EVs and NVEPs, we chose markers based on a literature survey. These included a membrane targeted Myristylation/Palmitoylation (MysP), indicative of membrane origin (15, 16), the canonical tetraspanin CD63 (17–19) recognized as a characteristic marker for EVs. ADP ribosylation factor 6 (Arf6) was also investigated due to its participation in shedding of lEVs, via the ectosomal route (20). Non-canonical markers for EVs were also studied including Rab GTPase 7a (Rab7a) (21, 22), and microtubule associated protein 1 light chain 3 beta (LC3b) (23–26), which have recently been shown to be involved in other potential EV release pathways. Finally, we assessed the capacity of NVEPs to mediate molecular cargo delivery to recipient cells, highlighting their potential as a next-generation therapeutic platform.

## Results

### ILVs’ Conserved Across Species: Meta-Analysis and In Situ Validation

To understand the maximum size of ILVs that can fit into MVB, we have performed a comprehensive analysis of published transmission electron microscopy (TEM) images. A total of 800 studies were identified using the PubMed search engine with the search query "(exosome AND TEM)" OR "(small+EV AND TEM)" OR "(MVB AND TEM)" OR "(ILV AND TEM)," conducted until June 22, 2025. We applied exclusion criteria to narrow down to those studies suitable for analysis. These exclusion criteria were (I) the absence of TEM micrographs of MVBs, (II) the absence of MVB TEM micrographs where ILVs were visible, (III) the absence of a scale bar, or (IV) if the TEM micrographs exhibited suboptimal resolution. Applying these criteria, 761 studies were excluded, resulting in 39 studies that featured well-defined TEM micrographs of MVBs with visible scale bars. These 39 studies were included in our analysis to estimate the diameter of ILVs within the MVBs (Fig. 1A). The selected studies were broadly categorized into six groups: Homo sapiens, Rodentia, Pisces, Plantae, Chlorophyta and other organisms. The category of other organisms was chosen for those species having a single study matching our inclusion criteria in this analysis. For the other categories, there are at least three studies which matched our inclusion criteria. For each study, the diameter of the ILVs within the MVBs was determined (representative measurements are shown in SI Appendix Fig. 1A). The measured diameter of each included study is shown. The cumulative mean ILV diameter observed across studies was 109.60 ± 36.77 nm (Fig. 1B). To validate the results from the literature, we performed TEM imaging on two different cell lines, human lung squamous cell carcinoma line (EBC-1) and human embryonic kidney cell line (HEK293), to measure ILV size experimentally (Fig. 1C). The mean ± standard deviation diameters observed for EBC-1 and HEK293 cells were 104.6 ± 12.53 nm and 106.7 ± 2.35 nm, respectively (Fig. 1D). These values are shown as ‘experimental group’ and are compared with the ‘literature group’ (Fig. 1E). There was no difference in the size of ILVs in the two groups. In summary, analysis of ILV size showed that they are approximately 100 nm in diameter with no examples surpassing 200 nm within 39 studies representing multiple species and kingdoms as well as our experimental work.

**Figure 1.**
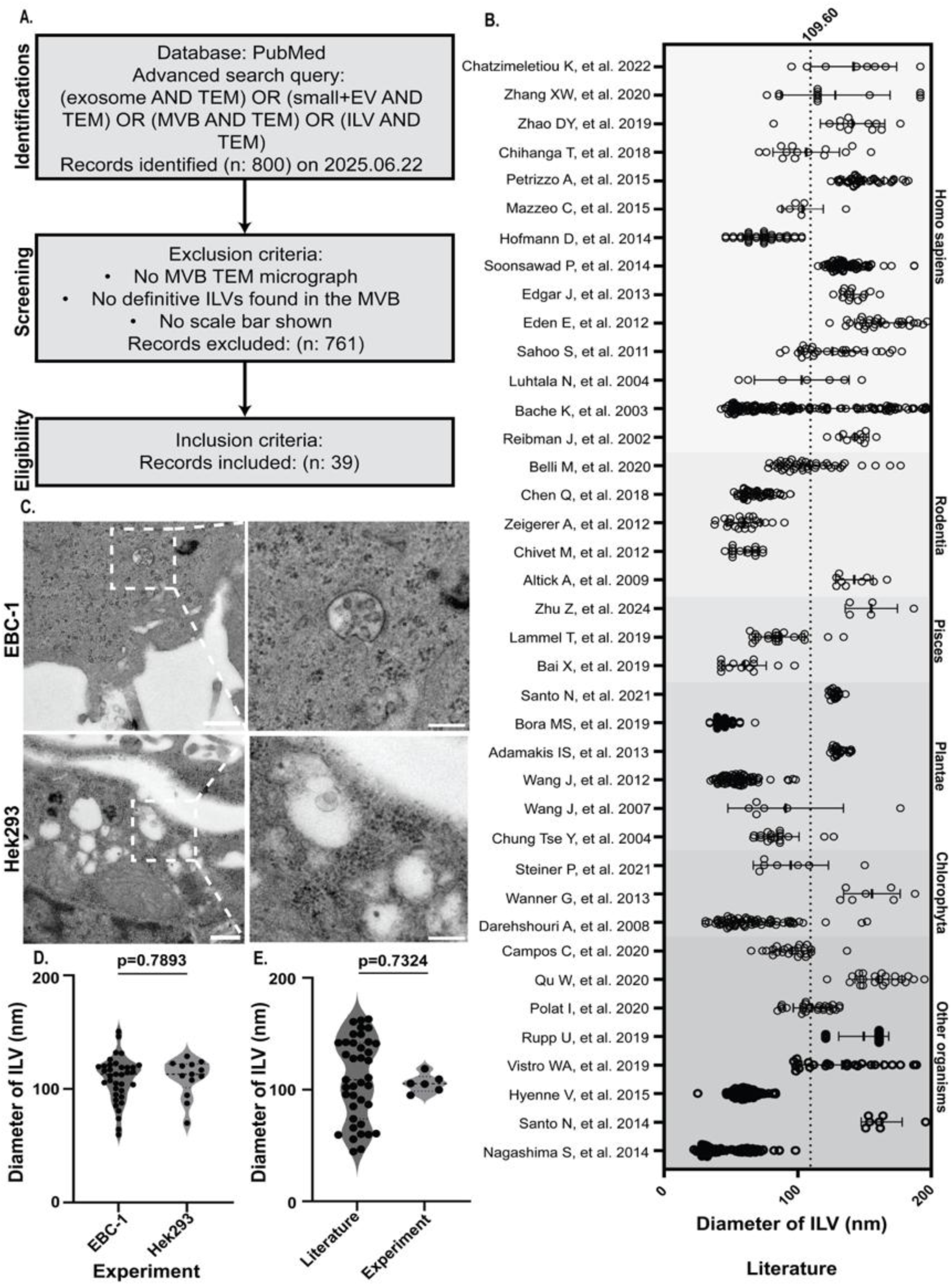
Comprehensive analysis of intraluminal vesicle (ILV) size distribution across species and experimental cell lines. (A) Summary of image analysis methodology based on a PubMed search, where 761 out of 800 studies were excluded due to not meeting inclusion criteria. (B) Distribution of ILV sizes within multivesicular bodies (MVBs) across kindgoms (Homo sapiens, Rodentia, Pisces, Plantae, Chlorophyta, and others), revealing a consistent diameter of less than 200 nm, with a mean of 109.60 ± 36.77 nm (termed “Literature”; n=39 studies). Data are presented as mean ± standard deviation. (C, D) Representative transmission electron microscopy (TEM) micrographs of MVBs in human lung squamous cell carcinoma cell line (EBC-1) (C) and Human Embryonic Kidney 293 cell line (HEK293) (D) cell lines, with measured ILV diameters of 104.60 ± 12.53 nm and 106.70 ± 2.35 nm, respectively (termed “Experiment”; data are mean ± standard deviation from three biological replicates). Scale bars: 500 nm (left panel) and 200 nm (right panel). (E) Statistical comparison showing no significant difference between ILV sizes from literature data and experimental measurements (p=0.7324, Mann-Whitney test).

### Comprehensive Characterization of EV Subpopulations and NVEPs Reveals Distinct Morphological and Marker-Protein Profiles

To understand the morphology and characteristics of EVs and NVEPs derived from HEK293 cells, we analyzed them with TEM which revealed their heterogenous morphology depending upon their size. EVs appeared spherical, cup-shaped, and varied in size, ranging from 41 to 849 nm with a visible lipid bilayer while 167k-NVEPs exhibited non-membranous structures (Fig. 2A. I-VI). Notably, 167k-NVEP exhibited to be significantly smaller size when compared to other EV subtypes (SI Appendix Fig. 1B). We determined the total concentration and size distribution of the 14k-lEVs and 100k-sEVs using nanoparticle tracking analysis (NTA). There was no significant difference in the concentrations of 14k-lEVs and 100k-sEV (Fig. 2B). The peak size analysis indicated that the 14k-lEV pellet had a median diameter of 190.80 ± 26.87 nm, while the 100k-sEV pellet had 125.20 ± 3.20 nm (Fig. 2C). Additionally, the zeta-potential was measured for 14k-lEVs and 100k-sEVs showing −28.07 ± 0.76 mV and - 18.13 ± 10.20 mV, respectively (SI Appendix Fig. 2A). The size and zeta-potential individual particle distributions are shown in SI Appendix Fig. 2B. When comparing the size of subpopulations of EVs and ILVs, 14k-lEVs were significantly larger and there was no significant size difference was observed between ILVs and 100k-sEV both (SI Appendix Fig. 1C, D).

**Figure 2.**
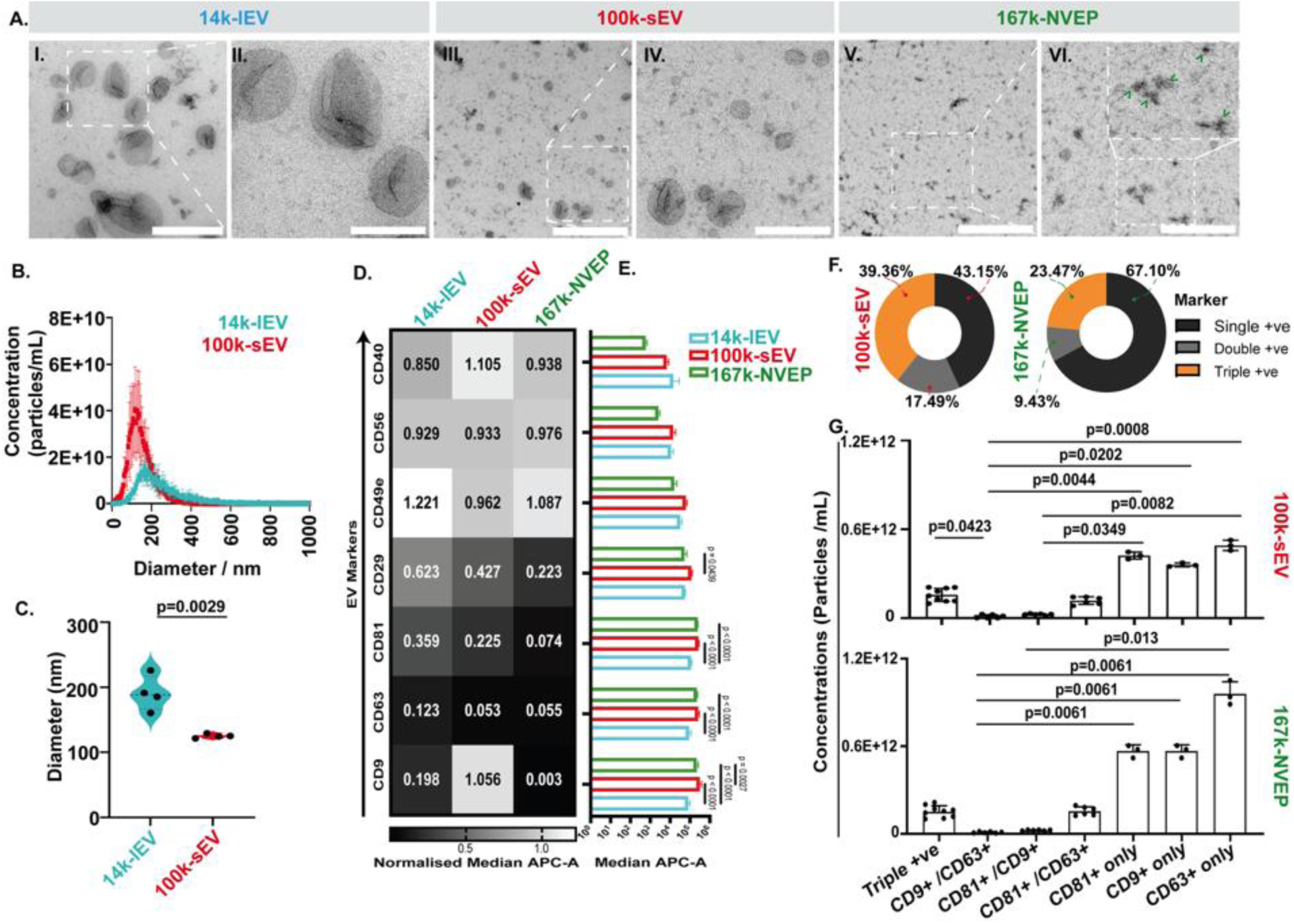
Characterization of size, surface charge, and marker enrichment in HEK293-derived EV subtypes and non-vesicular extracellular particles (NVEPs). (A) Transmission electron microscopy (TEM) images of 14k-IEV (complete field of view - I, magnified - II), 100k-sEV (complete field of view - III, magnified - IV), and 167k-NVEP (complete field of view - V, magnified - VI) fractions derived from HEK293 cells. 14k-IEV and 100k-sEV fractions exhibit clear lipid bilayer-enclosed structures, while 167k-NVEP predominantly contains non-vesicular particles (green arrows). Scale bars: 500 nm (left panels), 200 nm (right panels). Images are representative of three biological replicates. (B) Nanoparticle tracking analysis (NTA) showing total mean concentrations of 5.29E+11 particles/mL for 14k-IEV and 9.60E+11 particles/mL for 100k-sEV fractions (n=4 biological replicates). (C) Violin plot comparing vesicle sizes between 14k-IEV (mean ± standard deviation diameter = 190.80 ± 26.87 nm) and 100k-sEV (mean ± standard deviation diameter = 125.20 ± 3.20 nm) fractions, with a statistically significant difference observed (p=0.0029, unpaired t-test; n=4 biological replicates). (D) Heatmap showing normalized median APC-A fluorescence intensities for seven EV surface markers (CD40, CD56, CD49e, CD29, CD81, CD63, CD9) detected on 14k-IEV (cyan), 100k-sEV (red), and 167k-NVEP (green) fractions. Values represent normalized median APC-A signal for each marker and EV subtype. (E) Quantitative bar graphs of median APC-A fluorescence intensity for each marker across the three EV subtypes, with statistical significance indicated (p-values from ordinary one-way ANOVA with Tukey’s multiple comparisons test). (F) Single-particle interferometric reflectance imaging sensing (SP-IRIS) analysis of vesicles positive for single (CD9+, CD63+, or CD81+), double (CD9+/CD63+, CD81+/CD9+, or CD81+/CD63+), and triple (CD9+, CD63+, and CD81+) markers in 100k-sEV and 167k-NVEP fractions. (G) Quantitative analysis revealing that 100k-sEV exhibited significantly higher expression of individual markers compared to dual or triple markers (p<0.05, Kruskal-Wallis test). A similar pattern was observed for 167k-NVEP, with CD63+ single markers predominantly and significantly expressed (p<0.05, Kruskal-Wallis test). Data represent mean ± standard deviation from at least three independent biological replicates.

We assessed the surface marker composition of 14k-lEV, 100k-sEV, and 167k-NVEP using the MACSPlex Exosome Kit. Positive gating was established by comparing PE-A vs. APC-A signals against phosphate-buffered saline (PBS) controls, and markers with signal exceeding background were considered positive (SI Appendix Fig. 2C). For each marker, the signal was first corrected by subtracting background from PBS and the highest negative isotype control signal (REA or mIgG1), resulting in the raw adjusted intensity. Each marker’s signal was divided by the average intensity of CD9, CD63, and CD81 to account for variations in marker abundance. This analysis revealed subtype-specific enrichment patterns: CD49e was highest in 14k-lEV, CD40 and CD9 were enriched in 100k-sEV, and CD49e also showed elevated levels in 167k-NVEP (Fig. 2D). Supplementary analysis further showed that CD146 was highly enriched in 14k-lEV and 100k-sEV, while most other markers showed minimal signals across subtypes (SI Appendix Fig. 2D). The tetraspanins CD9, CD63, and CD81 had the highest absolute signal in 100k-sEV, consistent with their known enrichment in small EVs (Fig. 2E). In the supplementary dataset, CD146 and HLA-ABC were most abundant in 14k-lEV, while HLA-DRDPDQ showed the highest signal in 100k-sEV (SI Appendix Fig. 2E).

Next, we used single-particle interferometric reflectance imaging sensor (SP-IRIS) to measure marker expression analysis of 100k-sEV and 167k-NVEP. Varying concentrations of single-positive (CD63+, CD9+, CD81+), double-positive (CD81+/CD63+, CD81+/CD9+, CD63+/CD9+), and triple-positive (CD81+/CD63+/CD9+) markers were observed in both 100k-sEVs and 167k-NVEPs (SI Appendix Fig. 2F). Specifically, 100k-sEV showed 43.15% single positivity, 17.49% double positivity, and 39.36% triple positivity, while 167k-NVEPs had 67.1% single positivity, 9.43% double positivity, and 23.47% triple positivity (Fig. 2F). The concentration of 100k-sEVs expressing CD81+/CD63+ was significantly higher than that of 167k-NVEPs. Additionally, 100k-sEVs expressing both CD63+ and CD81+ were significantly more abundant than those expressing CD81+/CD9+. The concentration of 100k-sEVs expressing all three markers (CD63+, CD9+, CD81+) was also significantly higher than those expressing only CD63+/CD9+. The triple-positive EVs had a significantly higher concentration compared to the double-positive CD63+/CD9+ group. For 167k-NVEPs, the concentration of particles with all three markers were significantly higher than those expressing CD63+/CD9+, and the concentration of 100k-sEVs and 167k-NVEPs expressing CD63+ alone was significantly greater than those expressing CD81+/CD9+ (Fig. 2G).

### NVEPs Exhibit Elevated Protein-to-Lipid Ratios and Unique Raman Signatures

The biochemical and label-free characterization of 14k-lEVs, 100k-sEVs, and 167k-NVEPs revealed distinct parameters that may serve as effective methods for distinguishing 167k-NVEP from EV subpopulations. The total lipid and protein contents of EVs and NVEPs were quantified by estimating the concentration, which was normalized per million cells seeded. The 167k-NVEP subpopulation exhibited significantly higher protein-to-lipid ratio compared to 100k-sEVs and 14k-lEVs, and in 100k-sEVs compared to 14k-lEVs (Fig. 3A). Raman spectroscopy was employed to distinguish the biochemical makeup between 100k-sEVs and 167k-NVEPs. While the overall spectral profiles of the 100k-sEV and 167k-NVEP populations appeared comparable, notable variations in peak intensities were observed. The Raman shifts recorded spanned from 500 to 2000 cm⁻¹ (Fig. 3B), with comprehensive peak assignments provided in the supplementary information (SI Appendix Fig. 3A). To further distinguish the biochemical profiles, principal component analysis (PCA) was performed. Examination of the Raman spectra within the 500–2000 cm⁻¹ range revealed that the primary sources of variance were in the 924–1003 cm⁻¹ and 1274– 1571 cm⁻¹ regions, corresponding to proteins, lipids, and nucleic acids, respectively. The main loading in principal component (PC)-1, which accounted for 14.1% of the variance and was the key factor differentiating 100k-sEVs from 167k-NVEPs, was detected at 1006 (assigned to phenylalanine, Phe), 1661 (assigned as Amide I), and 1441 cm⁻¹ (assigned as saturated lipid), corresponding to proteins and lipids. A 3D PCA score plot constructed using PC1 (14.1%), PC2 (13.8%), and PC3 (10.6%), exhibited a 99% separation between the two populations (Fig. 3C). A heat map of the Max Height for specific Raman shifts, including 1006 cm⁻¹ (assigned to phenylalanine, Phe) and 1661 cm⁻¹ (assigned to Amide I, proteins), showed that these Raman fingerprints were predominantly enriched in 167k-NVEPs compared to 100k-sEVs (Fig. 3D). This enrichment indicates a biochemical distinction in protein content between the two populations, particularly for protein structures represented by phenylalanine side chains and the Amide I band. At 850 cm⁻¹, representing C-C bonds (assigned to the protein backbone), 100k-sEVs showed significantly higher levels compared to 167k-NVEPs (Fig. 3E). In contrast, Raman shifts at 1006 cm⁻¹ (assigned to phenylalanine, an aromatic residue of proteins) (Fig. 3F) and 1661 cm⁻¹ (assigned to amide I, proteins) were significantly higher in 167k-NVEP (Fig. 3G). No significant differences were observed at other Raman shifts (SI Appendix Fig. 3B). Notably, the analysis revealed that the 167k-NVEP population exhibited a significantly higher protein-to-lipid intensity ratio compared to 100k-sEV (Fig. 3H), suggesting that 167k-NVEP is characterized by an enrichment in proteins relative to lipids. This distinction underscores the unique biochemical properties of 167k-NVEPs in comparison to 100k-sEVs.

**Figure 3.**
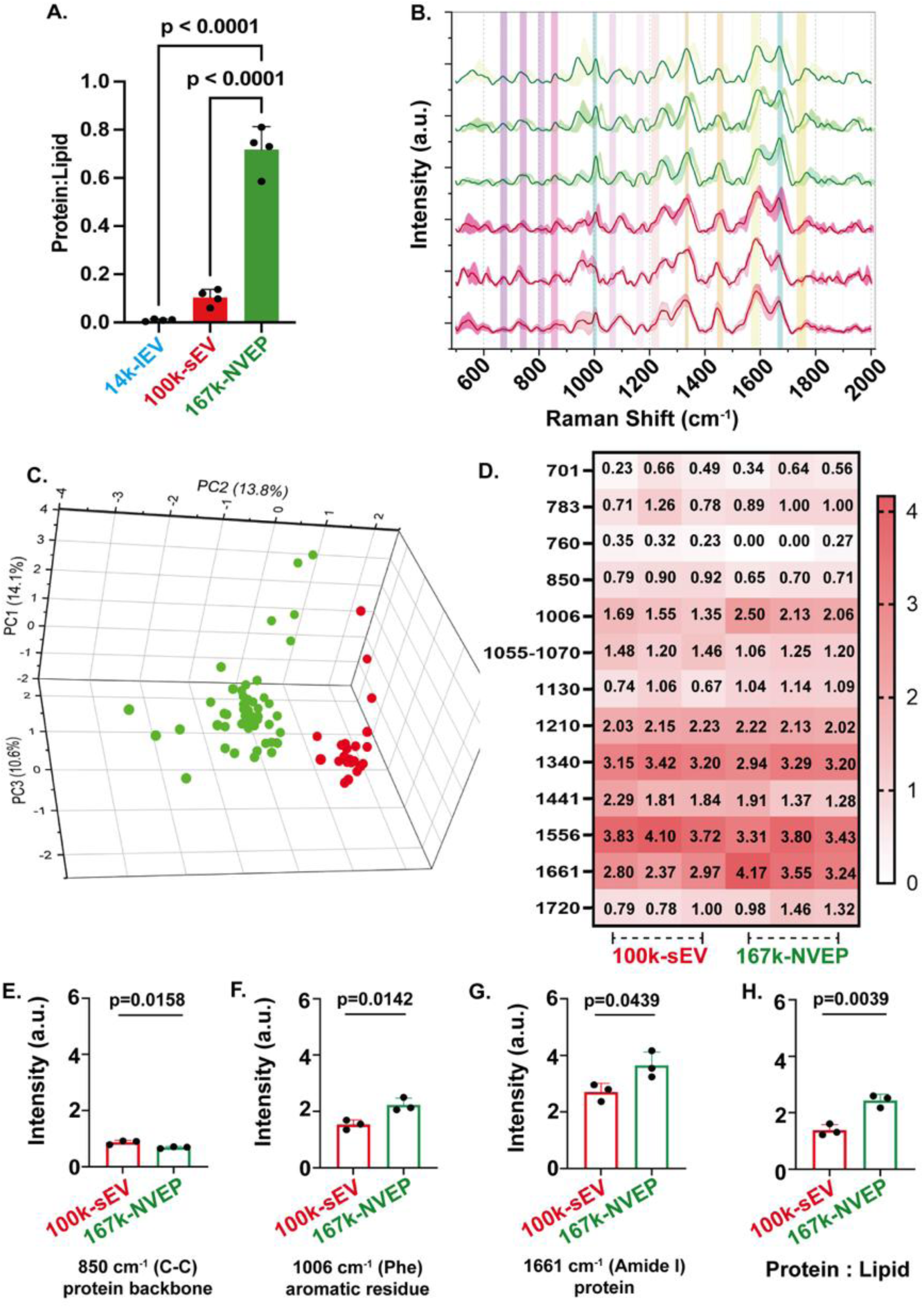
Biochemical characterization and Raman spectroscopy of EVs and non-vesicular extracellular particles (NVEPs). (A) Protein-to-lipid ratio is significantly higher in the 167k-NVEP fraction (0.72 ± 0.09) compared to both EV subtypes (p<0.0001, Ordinary One-way ANOVA with Tukey’s multiple comparisons test). Data are presented as mean ± standard deviation (n=4). (B) Raman spectroscopy of 100k-sEV (red) and 167k-NVEP (green) fractions, with solid lines representing average Raman spectra and shaded areas indicating standard deviation, showing shifts between 500 and 2000 cm^-1^ (n=3); the highlighted area are the representative Raman vibrational band assignments for HEK293-derived 100k-sEV and 167k-NVEP. (C) 3D scatter plot of Principal component analysis (PCA) showing cluster separation between 100k-sEV and 167k-NVEP fractions (n=3). (D) Heat map of Raman spectral deconvolution showing key peaks in 100k-sEV and 167k-NVEP fractions. (E) The 850 cm^-1^ band (C-C protein backbone, 0.87 ± 0.07 a.u) is predominant in 100k-sEV (p=0.0158, unpaired t test), (F) while the 1006 cm^-1^ (phenylalanine, 2.23 ± 0.23 a.u) and (G) 1661cm^-1^ (amide I, 3.65 ± 0.47 a.u) bands are enriched in 167k-NVEP (p=0.0142 and p=0.0439, respectively, unpaired t test). (H) Protein-to-lipid ratio in 167k-NVEP is significantly higher than in 100k-sEV (p=0.0039, unpaired t test), which was compared between 1661 cm^-1^ band (assigned as proteins) and 1441 cm^-1^ band (assigned as saturated lipids) of Raman Fingerprint. Data are presented as mean ± standard deviation (n=3).

### Lipidomic profiling reveals distinct lipid signatures in NVEPs

To characterize the lipid composition of 167k-NVEPs and compare it to 100k-sEV and secreting cells, we conducted lipidomic analysis using LION and LipidSig 2.0 (27–30). Hierarchical clustering heat maps which include lipid classification, chemical and physical properties (fatty acid length and unsaturation, headgroup charge, intrinsic curvature, membrane fluidity, bilayer thickness), function, and sub-cellular component (predominant sub-cellular localization) illustrate significant lipidomic variations between 100k-sEVs and 167k-NVEPs (Fig. 4A, B). 100k-sEV exhibited higher concentrations of sphingomyelins (SM), hexosylceramides (HexCer), and unsaturated fatty acids, while 167k-NVEP showed significant upregulated of ST0102 represents steryl esters (Fig. 4A). Secondly, to infer the potential subcellular origin of the lipid cargo in 100k-sEVs and 167k-NVEPs, we performed compartment-based lipid enrichment analysis. This revealed that 167k-NVEPs were enriched in lipid species typically associated with the endosome–lysosome system and lipid droplets, whereas 100k-sEVs showed higher abundance of lipids linked to the Golgi apparatus and plasma membrane (Fig. 4B). To examine how lipids are differentially distributed from the secreting cell into specific EV subtypes, we conducted PCA on the lipidomic profiles of parent HEK293 cells, 100k-sEVs, and 167k-NVEPs. PCA demonstrated clear separation of the three sample types along PC1 (74.4%), with the 100k-sEVs and the 167k-NVEPs grouping separately from cells (Fig. 4c). This distinction is also evident in the PCA of specific EV-related lipid classes, including free cholesterol (FC), SM, phosphatidylethanolamine plasmalogen (PE-P), phosphatidylethanolamine plasmalogen (PE-P), cholesteryl esters (CE), phosphatidylcholine (PC), phosphatidylethanolamine (PE), phosphatidylinositol (PI), and triglycerides (TG) (SI Appendix Fig. 4A). Quantification of lipid species showed CE predominance in 167k-NVEP compared to 100k-sEV or secreted cells, while 100k-sEV was more abundant with the FC, SM, PE-P whereas cells were more abundant with PC, PE, PI, TG, (Fig. 4D) whereas other lipid species found to be mostly enriched in the secreting cell except in 100k-sEVs, which showed notably higher levels of plasmalogen phosphatidylcholine (PC-O) and phosphatidic acid (PA), both essential for membrane dynamics and vesicle formation (SI Appendix Fig. 4B). These findings align with previous research indicating that EVs possess distinct lipid profiles, which can impact their structural characteristics and functional roles (31, 32). LipidSig 2.0-based enrichment analysis confirmed compositional distinctions: CE was elevated in 167k-NVEPs (Fig. 4e) while the volcano plot shows triglyceride 48:4 and CE species (20:4, 16:1, 16:0, 14:0) as NVEP-enriched. Conversely, the top five lipids enriched in 100k-sEVs were diacylglycerol 36:1, PI 34:1, PE 32:1, phosphatidylserine (PS) 40:5, and phosphatidylglycerol (PG) 34:1 (Fig. 4F). Notably, carbon chain length distributions diverged significantly: 167k-NVEPs exhibited higher abundance of 16-, 18-, and 20-carbon chains, while 100k-sEVs were enriched in 32-, 34-, 36-, and 42-carbon chains (Fig. 4G).

**Figure 4.**
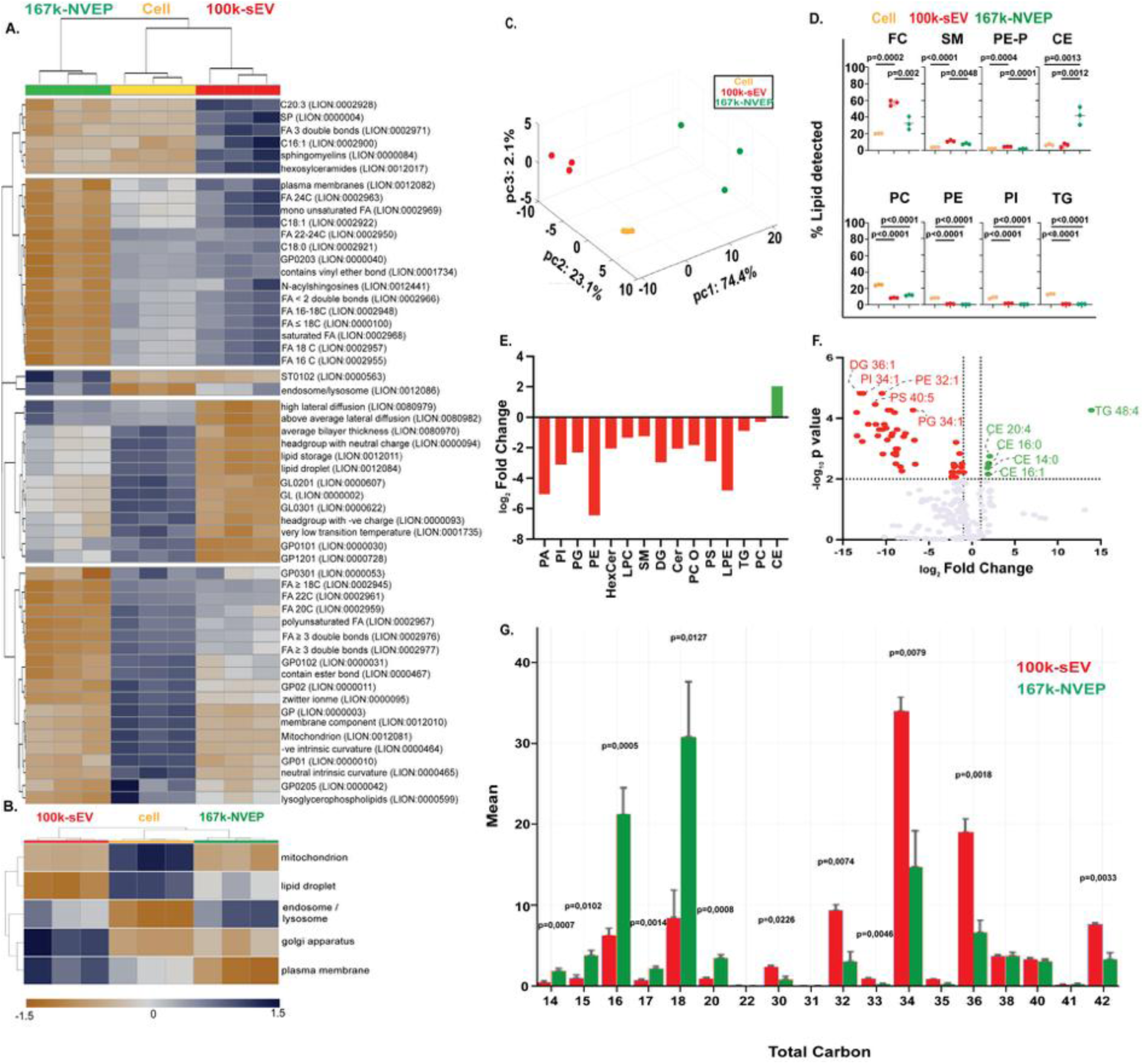
Lipidomic profiling and LION-term enrichment analysis of HEK293 cells, 100k-sEV, and 167k-NVEP. (A) Heatmap of scaled lipid amounts (z-scores: yellow, < 0; blue, > 0) from subcellular lipidomic data for cells, 100k-sEV, and 167k-NVEP. Lipids were clustered into five groups using hierarchical clustering. (B) Heatmap based on Lipid Ontology (LION)-term predictions, showing associations of lipid species with cellular compartments via hierarchical clustering. Lipids in 167k-NVEP are primarily linked to endosome/lysosome compartments, while 100k-sEV lipids are associated with endosome/lysosome, Golgi apparatus, and plasma membrane. (C) Mass spectrometry analysis showing the percentage distribution of lipid species. 100k-sEV is predominantly enriched in free cholesterol (FC), sphingomyelin (SM), and phosphatidylethanolamine-based plasmalogen (PE-P), whereas 167k-NVEP is highly enriched in cholesteryl ester (CE). Cells are enriched in phosphatidylcholine (PC), phosphatidylethanolamine (PE), phosphatidylinositol (PI), and triglycerides (TG). (D) Three-dimensional principal component analysis (PCA) plot displaying the distribution of cells (yellow), 100k-sEV (red), and 167k-NVEP (green) based on molecular signatures. Each point represents an independent biological replicate (n=3). Axes correspond to principal components 1, 2, and 3, explaining 74.4%, 23.1%, and 2.1% of total variance, respectively, demonstrating distinct clustering of each group. (E) Bar graph illustrating log₂ fold change in abundance of major lipid classes between 100k-sEV (red) and 167k-NVEP (green). Most lipid species, including phosphatidic acid (PA), PI, phosphatidylglycerol (PG), PE, hexosylceramide (HexCer), lysophosphatidylcholine (LPC), SM, diacylglycerol (DG), ceramide (Cer), PC, phosphatidylserine (PS), lysophosphatidylethanolamine (LPE), and TG, are significantly depleted in 100k-sEV relative to 167k-NVEP, while CE is enriched in 167k-NVEP. (F) Volcano plot depicting log₂ fold change (x-axis) and statistical significance (-log_10_(p value), y-axis) for individual lipid species comparing 100k-sEV and 167k-NVEP. Lipids significantly enriched in 100k-sEV (red) include DG 36:1, PI 34:1, PE 32:1, PS 40:5, and PG 34:1. Lipids significantly enriched in 167k-NVEP (green) include TG 48:4 and CE (CE 20:4, CE 16:0, CE 14:0, CE 16:1). Dotted lines indicate thresholds for statistical significance (p<0.01) and fold change. All data represents mean values from three independent biological replicates (n=3). (G) Carbon chain length distribution in 100k-sEV and 167k-NVEP fractions analyzed via LipidSig differential expression. The 167k-NVEP fraction (green bars) shows significant enrichment in lower carbon chain lipids (C16-C20, p<0.001 at C16 and C20), while 100k-sEV (red bars) exhibits selective enrichment at longer chain lengths (C31, C34, C36, p<0.01). Data represent mean relative abundance ± standard deviation from n=3 biological replicates, with statistical significance determined by unpaired t-tests (p<0.001, p<0.01). Findings suggest distinct membrane properties between EV subtypes and 167k-NVEPs.

### Subcellular Localization of Molecules Implicated in EV biogenesis and Comparative Analysis of Membrane Association

To trace the biogenesis route of the EV subtypes and 167k-NVEP, we expressed in HEK293 cells various eGFP-tagged markers (Arf6, CD63, LC3b, MysP, Rab7a) and developed a quantitative membrane association assay. Confocal micrographs were segmented to define nuclei and plasma membranes. For each eGFP signal, we calculated the minimum distance to nuclear and plasma membranes, deriving a membrane association percentage (Fig. 5A; methodology schematic and representative images in Fig. 5B) and the workflow pipeline was shown by schematic representation in SI Appendix Fig. 8. This approach allowed us to quantitatively assess the extent of membrane association for each marker in comparison to MysP. Analysis of confocal images from cells expressing these markers (n=5) indicated that Rab7a, and Arf6 showed significantly lower membrane association than MysP (Fig. 5C). The reduced membrane association of Rab7a, and Arf6 implies that these markers are primarily located within endosomal compartments rather than at the plasma membrane, which is consistent with their known functions in ILV formation and endocytosis regulation (33). Autophagosome marker LC3b exhibited levels of membrane association comparable to MysP, with no statistically significant difference noted. This localization aligns with previous studies describing the interaction of LC3b with membrane-associated autophagic structures (34). Furthermore, the tetraspanin protein CD63, known for its role in exosome biogenesis, did not show significant differences in membrane association compared to MysP, suggesting a distribution across both endosomal and plasma membrane locations. In contrast, Arf6, which is also associated with vesicle trafficking closer to the nucleus, displayed a distinct pattern in this study. This observation is in line with earlier reports highlighting the role of Arf6 in nuclear-proximal vesicle transport (33).

**Figure 5.**
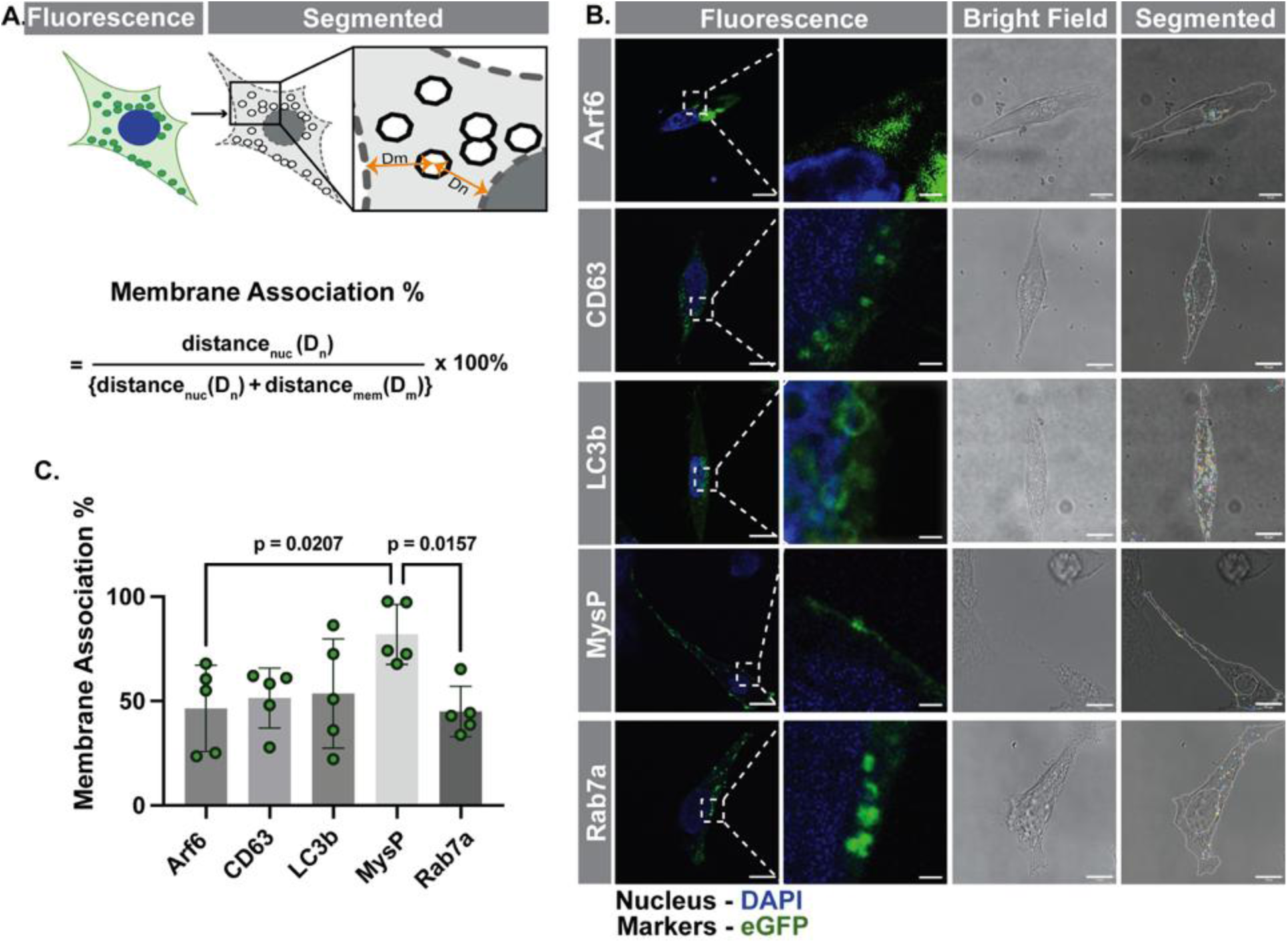
Subcellular localization and membrane association of EV biogenesis markers in HEK293 cells. (A) Schematic representation of the sub-cellular localization of the over expressed marker including ADP ribosylation factor 6 (Arf6), CD63, microtubule associated protein 1 light chain 3 beta (LC3b), Myristylation/Palmitoylation (MysP) and Rab GTPase 7a (Rab7a); tagged with eGFP over expressing in single HEK293 cells was visualized under confocal microscope. From the fluorescence image and bright field confocal micrographs, nucleus, plasma membrane and eGFP signal were segmented. Then we calculated the shortest distance between eGFP segments to plasma membrane and nucleus, with which we derive the Membrane association percent directly proportional to the association of close to the plasma membrane. (B) Representative confocal micrographs of eGFP-tagged markers associated with extracellular vesicle (EV) biogenesis (CD63, Rab7a, Arf6), autophagosomes (LC3b), and a lipidated membrane marker (MysP), individually transfected into HEK293 cells. Images show fluorescence (left), brightfield (centre), and segmented (right) views to assess membrane association, compiled from five biological replicates. Scale bar, 10 μm and 1 μm (fluorescence right). (C) Comparative analysis of membrane association percentages, using MysP (81.91 ± 14.40%) as a benchmark. No significant differences were observed for CD63 (51.46 ± 14.36%), LC3b (53.62 ± 26.13%) when compared to MysP (p>0.05). Significant differences were detected for Arf6 (46.46 ± 20.70%, p=0.0250) and Rab7a (45.01 ± 12.07%, p=0.0184) relative to MysP, as determined by ordinary one-way ANOVA with Dunnett’s multiple comparisons test. Data are presented as mean ± standard deviation from five independent experiments (n=5).

### Relative marker distribution Distinguishes EV Subtypes and NVEPs Across Platforms

EV subtypes and 167k-NVEPs were isolated from HEK293 cells expressing selected markers for characterization as well as from non-transfected (NT) control conditions. NTA was used to determine concentration and size of 14k-lEV and 100k-sEVs (SI Appendix Fig. 5A-D). Total lipid content was significantly higher in 14k-lEVs than in 100k-sEVs or 167k-NVEPs (SI Appendix Fig. 5E). Conversely, 167k-NVEPs exhibited significantly higher protein content than either 14k-lEVs or 100k-sEVs (SI Appendix Fig. 5F). Critically, the protein-to-lipid ratio of 167k-NVEPs was significantly higher than that of all EV subtypes, consistent across both NT and overexpressing conditions (SI Appendix Fig. 5G). Following the Minimum Information for Flow Cytometry studies of Extracellular Vesicles (MIFlowCyt-EV) guideline (35), we estimated the eGFP% and concentration (events/mL) of the over-expressed EV subtypes, where gating was made in respect to their NT counter parts (SI Appendix Fig. 6A). Histogram showing the comparison between the marker expressing EV subtypes and their NT counterparts (SI Appendix Fig. 6B). No markers showed any significant difference in the eGFP% between their EV subtypes (SI Appendix Fig. 6C), whereas LC3b and MysP expressing 14klEV events/mL were significantly higher than their 100k-sEV counterparts (SI Appendix Fig. 6D). Conversely, Rab7a expressing 100k-sEV events/mL was significantly higher than 14k-lEV (SI Appendix Fig. 6D). Taken together, we found that no marker is exclusive in either EV subtypes, but it is possible to see an enrichment in CD63 and MysP for 14k-lEV or CD63 and Rab7a for 100k-sEV (SI Appendix Fig. 6E).

**Figure 6.**
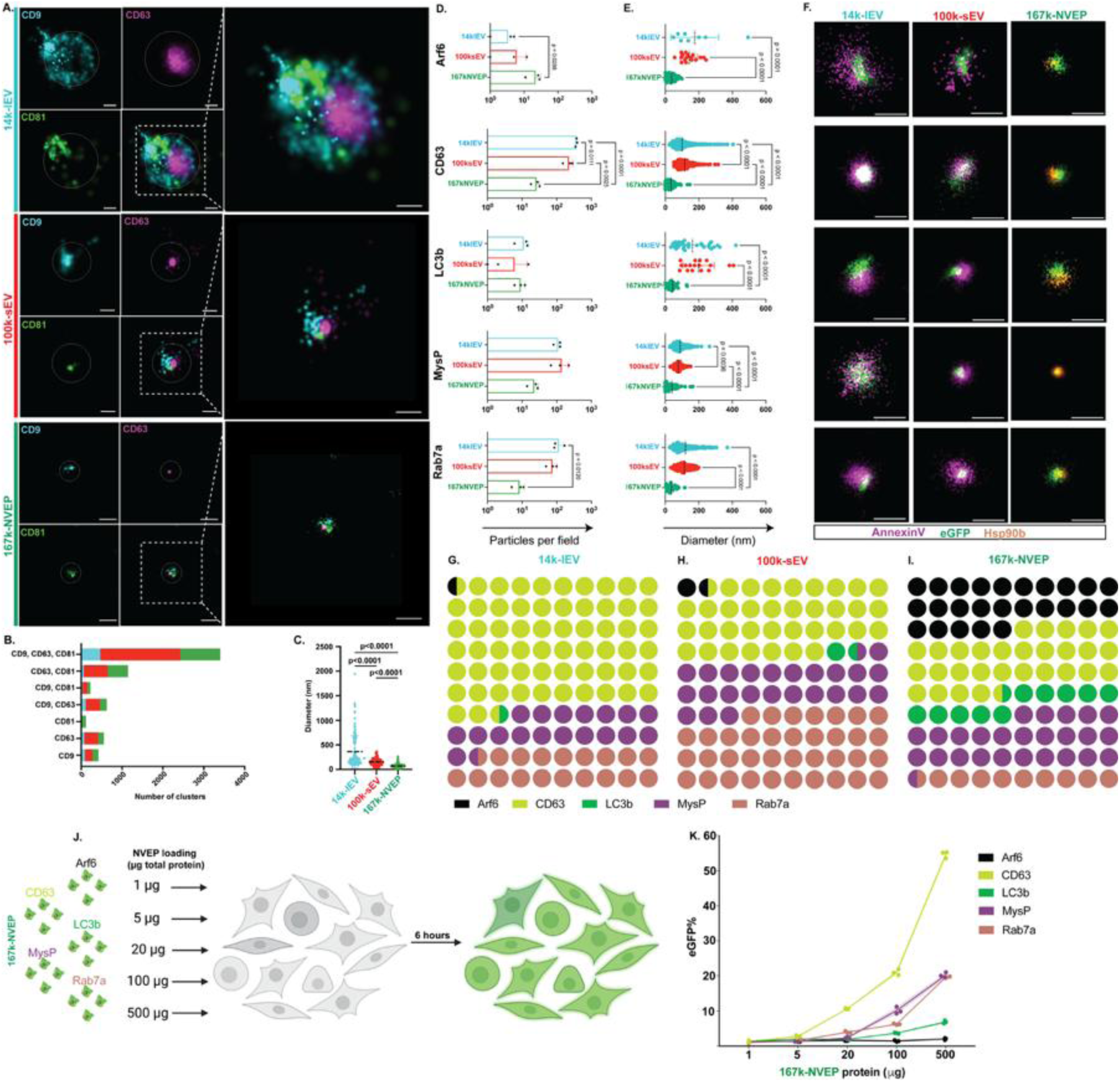
Super-resolution profiling and uptake dynamics of native EVs and 167k-NVEPs reveal subtype-specific marker landscapes and functional heterogeneity. (A) Super-resolution characterization of native EVs and 167k-NVEPs by direct Stochastic Optical Reconstruction Microscopy (dSTORM). Representative dSTORM images of immunolabeled CD9, CD63, and CD81 were acquired for both native EVs and NVEPs. Particles were stained with fluorophore-conjugated antibodies against the tetraspanins CD9, CD63, and CD81. Scale bars, 100 nm. (B) Quantification of cluster compositions showing the number of single-, double-, and triple-positive tetraspanin clusters in 14k-lEV (cyan), 100k-sEV (red), and 167k-NVEP (green). Statistical analysis was performed using Ordinary two-way ANOVA. (C) Scatter plot displaying the diameter distribution of individual particles across 14k-lEV, 100k-sEV, and 167k-NVEP populations. Mean values are indicated by horizontal dashed lines (p<0.0001, Kruskal–Wallis test). (D) Bar plots represent the number of single EVs and NVEPs per field expressing the indicated markers—Arf6, CD63, LC3b, MysP, and Rab7a —detected in 14k-lEVs (cyan), 100k-sEVs (red), and 167k-NEVPs (green). Across biological replicates (n=3), 167k-NEVPs exhibited significantly higher numbers of Arf6-positive particles per field compared to 14k-lEVs (p=0.0288, Ordinary one-way ANOVA with Tukey’s multiple comparisons test). Similarly, significant differences were observed in the expression patterns of other markers, including higher CD63 compared to 100ksEV (p=0.0111, Ordinary one-way ANOVA with Tukey’s multiple comparisons test) and compared to 167k-NEVPs (p=0.0001, Ordinary one-way ANOVA with Tukey’s multiple comparisons test) and significant increase of Rab7a expressing 14k-lEVs compared to 167k-NEVPs (p = 0.0102, Ordinary one-way ANOVA with Tukey’s multiple comparisons test). (E) Diameter measurements of individual particles show that 14k-lEVs were significantly larger than both 100k-sEVs and 167k-NEVPs, and that 100k-sEVs were significantly larger than 167k-NEVPs. (F) Representative super-resolution images of single EV and NVEP particles with their respective markers. 14k-lEVs and 100k-sEVs were labeled with Annexin V (magenta) and anti-EGFP antibody (green), while 167k-NEVPs were stained with anti-HSP90β (orange) and anti-EGFP (green). Scale bar: 100 nm. (G-I) Parts of whole matrix plots display the mean number of particles per field positive for each subcellular marker in 14k-lEVs (G), 100k-sEVs (H), and 167k-NEVPs (I), where the markers include Arf6 (black), CD63 (chartreuse), LC3b (green), MysP (purple), and Rab7a (coral). Number of dots reflects the average number of positive particles per field. Notably, CD63 (29.77%), Arf6 (24.81%), MysP (25.57%) were enriched in 167k-NEVPs; Rab7a (37.15%) and CD63 (35.38%) in 100k-sEVs; and CD63 (61.89%), Rab7a (18.74%) and MysP (18.08%) in 14k-lEVs. While several markers showed subtype-specific enrichment, none were exclusive to a particular EV class. Rather, it is the relative enrichment pattern that distinguishes each vesicle population. (J) Schematic of NVEP uptake assay. HEK293 cells were incubated with increasing doses (1, 5, 20, 100, and 500 μg total protein) of 167k-NVEPs expressing different EGFP-tagged protein markers (Arf6, CD63, LC3b, MysP, Rab7a). eGFP-positive cells were quantified via flow cytometry. (k) Dose-dependent uptake efficiency of 167k-NVEPs by marker type. Quantification of eGFP-positive cells showed that all tested markers facilitated dose-responsive uptake. CD63-expressing 167k-NVEPs yielded the highest cellular uptake (54.60 ± 0.95% at 500 μg). Data represent mean ± standard deviation from three independent flow cytometric experiments (n=3).

In addition to these measurements, we employed direct stochastic optical reconstruction microscopy (dSTORM) to conduct single-particle analysis of 14k-lEVs, 100k-sEVs, and 167k-NVEPs. Firstly, we checked the distribution of the known canonical markers of the EVs including CD9, CD63 and CD81 across NT EV subtypes and 167k-NVEP (Fig. 6A), where we observed that 100k-sEV triple positivity clusters were significantly higher than 14k-lEV and 167k-NVEP (Fig. 6B). In terms of size, 14klEV was larger followed by 100k-sEV and 167k-NVEP (Fig. 6C). For the marker expressing EV subtypes, we used Annexin V as a membrane marker due to its phosphatidylserine-binding capability, along with an anti-GFP antibody targeting eGFP-tagged markers previously studied. For 167k-NVEPs, which lack a membrane for Annexin V binding, we utilized the reported marker HSP90ß (8, 36), as 167k-NVEP single positive annexin-V clusters was significantly lower than 100k-sEV (SI Appendix Fig. 9A). A representative image shows the cluster finding by applying the EV profiling in the CODI webtool (SI Appendix Fig. 9B).

We evaluated identified diameters for all markers expressing EV subtypes and 167k-NVEP samples expressing the selected markers, comparing these parameters across different groups. CD63-expressing 14k-lEVs and 100k-sEVs had significantly higher particle counts than 167k-NVEPs, while Arf6-expressing 167k-NVEPs exceeded 14k-lEVs in particles per field (Fig. 6D). The diameter measurements of 14k-lEVs were consistently larger than those of 100k-sEVs and 167k-NVEPs (Fig. 6E). Representative EVs and NVEPs for all the markers used in this study were shown in Fig. 6f. Our data revealed distinct marker abundances across subpopulations. None of the markers used in this study seem to be exclusive for any EV subtypes or 167k-NVEPs. Interestingly, 14k-lEV was more abundant with CD63, MysP (Fig. 6g); 100k-sEV was more abundant with CD63, Rab7a (Fig. 6h) and 167k-NVEP was more abundant with Arf6 and CD63 (Fig. 6i). To assess the consistency of size measurements across orthogonal platforms, we compared the particle diameters obtained from dSTORM with those measured by NTA. Many diameters were consistent across platforms except for CD63, MysP and Rab7a expressing 14k-lEV as well as CD63 and MysP expressing 100k-sEV being significantly larger by NTA than dSTORM (SI Appendix Fig. 7A, B).

Finally, we assessed if 167k-NVEPs can deliver eGFP to recipient cells. Different protein amounts (1 μg, 5 μg, 20 μg, 100 μg, 500 μg) were added to native HEK293 cells (Fig. 6J), where we observed that CD63 expressing 167k-NVEPs had the highest efficiency of eGFP transfer reaching 54.60% of treated cells becoming eGFP positive (Fig. 6K).

## Discussion

Our integrated approach—combining cross-kingdom meta-analysis, multiparametric characterization including biochemical profiling, Raman spectroscopy, lipidomic profiling, and advanced imaging—reveals fundamental principles governing EV heterogeneity and identifies NVEPs as sterol-rich entities with unique biogenetic origins. The remarkable conservation of ILV diameters—centering on ∼109 nm and not exceeding 200 nm—holds true not only in mammalian systems but also across phylogenetically distant lineages (animals, plants, fungi, and protists), as revealed by our meta-analysis of published TEM data and corroborated by independent measurements. This cross-kingdom consistency reinforces the MISEV2023 guideline (7), which defines exosome-sized vesicles as ≤200 nm, which is the fate of ILVs within MVBs. Nonetheless, several considerations must be taken before labelling all EVs under 200nm as exosomes. First, TEM size estimates are sensitive to sample preparation artifacts (fixation, dehydration, staining) that can compress or inflate vesicles. Second, ILVs measured *in situ* before secretion may not reflect their final size in fluids since after secretion other events could occur such as small EVs of exosomal origin fusing to make larger ones or large EVs of ectosomal origin breaking up into smaller ones. In such potential scenarios, it would still hold true that under the current biogenesis classification accepted for EVs, exosomes cannot exceed 200nm in size but new models would be needed to properly reflect a more complex and dynamic reality that our current classification does not reflect.

Here in this study, we have used HEK293 cell-derived EVs and NVEPs. This is due to the unique advantages of HEK293 for EV and NVEP research, as substantiated by leading studies in the field (13, 37, 38). HEK293 cells are human-derived, highly amenable to genetic engineering, and capable of robust, scalable, and serum-free EV production—enabling precise manipulation of EV surface features and cargo for mechanistic studies and therapeutic development. These attributes ensure consistent particle composition and purity and minimize batch variability.

To have better understanding the NVEPs composition and source of origin, we employed orthogonal techniques to compare EV subpopulations to NVEPs including TEM, NTA, zeta potential measurement, multiplexed-bead assay and high-resolution flow cytometry, SP-IRIS, biochemical assays, Raman spectroscopy, lipidomic, and super-resolution microscopy. TEM images confirmed the presence and absence of lipid bilayer membranes in EV subpopulations (14k-lEV and 100k-sEV) and 167k-NVEP (8), respectively. This morphological contrast aligns with previous reports describing exomeres, a type of NVEP, as non-membranous entities (8). Overall distribution of the canonical EV markers was found to be localized in the NVEPs populations. Interestingly, we observed that NVEP exhibited high protein to lipid ratio, which can readily distinguish and characterize through this straightforward, reliable, reproducible, and cost-effective method. Characteristic bands corresponding to major EV components were observed in the fingerprint region of Raman spectra for all EV samples, consistent with previous findings (39, 40). Our Raman spectroscopy results showed a significantly high protein-to-lipid ratio of 167k-NVEP, aligning with our biochemical analysis data. Additional Raman spectroscopy should be conducted to better understand the C-H Stretching (2800-3000 cm⁻¹) region, which can be further correlated with lipidomic data to elucidate characteristic differences between 167k-NVEPs and 100k-sEVs.

Lipidomic profiling revealed 167k-NVEPs as a biochemically distinct extracellular nanoparticle population, enriched in steryl esters and CE compared to 100k-sEVs (41, 42). The lipidomic analysis reveals that 167k-NVEPs possess a lipid composition sharply distinct from both 100k-sEVs and their parent cells, underscoring NVEPs as a discrete class of extracellular particles with unique biophysical properties and likely specialized functions. 100k-sEV were found to contain lipids associated with intercellular communication, though certain enriched lipid species suggest the inclusion of additional structures, potentially from amphisome (43). The 100k-sEV fraction also showed an abundance of sphingolipids, including sphingomyelins (SM) and hexosylceramides (HexCer), which are essential for exosome formation and play roles in brain cell function (44, 45). A previous study showed that SM and HexCer which are enriched in 100k-sEV can support membrane structure and promote amyloid-β clearance through exosome secretion in microglia, suggesting neuroprotective roles for these EVs (46). NVEPs were markedly enriched in lipid species that preferentially localize to endosomal/lysosomal membranes and lipid droplets—whereas sEVs contained higher levels of SM, HexCer, and polyunsaturated fatty acids. The prevalence of esterified sterols implies altered membrane rigidity and fusion dynamics, raising the possibility that NVEPs serve specialized functions in hydrophobic cargo transport or modulation of recipient cell lipid metabolism. This distinct lipid signature suggests a non-endosomal, possibly secretory lysosomal origin, consistent with previous observations that exomeres may arise via alternative non-canonical secretion pathways (47, 48).

To elucidate the molecular determinants of NVEP biogenesis, we interrogated the subcellular localization of eGFP-tagged markers in HEK293 cells by confocal microscopy coupled with an in-house developed membrane association assay. Our analyses revealed that Arf6 and Rab7a exhibited perinuclear localization distinct from canonical endolysosomal markers CD63 and LC3b, as compared to a membrane-targeted control. Super-resolution dSTORM imaging confirmed the presence of classical tetraspanin markers—CD63, CD9, and CD81—on all extracellular vesicle (EV) subtypes. However, these markers were significantly less abundant on 167k-NVEPs, suggesting a non-vesicular molecular composition. Similarly, Annexin V binding revealed much lower phosphatidylserine exposure in 167k-NVEPs, consistent with the lack of a typical bilayer membrane and the asymmetrical phospholipid distribution normally seen in EVs. We observed that no single marker—including Arf6, CD63, LC3b, MysP, and Rab7a—was uniquely restricted to a specific EV or NVEP subtypes, indicating that there is no marker exclusivity for the subtype but rather a heterogeneously distributed marker landscape. Finally, dose-dependent uptake assays demonstrated that CD63-eGFP enriched NVEPs loaded at 500 µg total protein delivered eGFP to HEK293 recipient cells, underscoring NVEPs’ potential as non-membranous therapeutic vehicles.

## Conclusions

Our cross-kingdom survey shows that intraluminal vesicles do not exceed ∼200 nm, which aligns with established exosome size criteria. We show that NVEPs display a distinct biochemical and molecular signature, notably an elevated protein-to-lipid ratio, as evidenced by both biochemical assays and Raman spectroscopy. Lipidomic profiling further indicates enrichment of CE and TG in NVEPs, implicating lipid droplets and endo/lysosomal membranes as probable sources. Furthermore, NVEPs can be used to deliver cargo to recipient cells, making them a promising, non-membranous platform for next-generation therapeutic delivery. Collectively, these findings suggest that NVEPs may form a compositionally and biogenetically distinct class of extracellular particles, warranting further investigation into their formation mechanisms and functional roles.

## Materials and Methods

### 1. Comprehensive image analysis of ILV

To conduct a comprehensive image analysis on the size of intraluminal vesicles (ILVs), we focused exclusively on studies indexed in PubMed. A literature search was conducted on 22 June, 2025, using the query: (exosome AND TEM) OR (small+EV AND TEM) OR (MVB AND TEM) OR (ILV AND TEM). This search yielded 800 records. After applying exclusion criteria—specifically, the absence of multivesicular body (MVB) transmission electron microscopy (TEM) micrographs, lack of definitive ILV measurements, or missing scale bars—761 original research articles were excluded. Ultimately, 39 original articles met the inclusion criteria and were included in this study.

### 2. Cell Culture

In this study, two derivatives of human embryonic kidney cells (HEK293) were utilized: HEK293T and 293A cell lines. HEK293T cells were obtained from the European Collection of Authenticated Cell Cultures (ECACC). 293A cell line was provided by Várnai Péter (Semmelweis University, Budapest, Hungary). Additionally, EBC-1 cell line was provided by Angéla Takács (Semmelweis University, Budapest, Hungary). All cells were cultured in Dulbecco’s Modified Eagle’s Medium (Catalogue # D6046; Merck Group, MA, USA) supplemented with 10% fetal bovine serum (Catalogue # 10270106; Gibco, NY, USA) and 1% penicillin–streptomycin (Catalogue # P4333; Merck Group, MA, USA). All cells were incubated at 37°C in a 5% CO_2_ environment and ≥ 90% humidity. Cells were passaged three times per week. Polymerase chain reaction (PCR) was routinely employed to assess cell cultures for mycoplasma contamination. The PCR primers utilized in this process were: GTTTGATCCTGGCTCAGGAYDAAC and GAAAGGAGGTRWTCCAYCCSCAC (49)

### 3. Molecular Cloning

In this study, we have used 2 recombinant plasmids, which were developed in-house. They are Myc_EGFP_Arf6_HA (SGb42) and MysP-eGFP (BMKb9). We have also used pEGFP-LC3 (addgene: Plasmid #24920, MA, USA), GFP-rab7 WT (addgene: Plasmid #12605, MA, USA), and CD63-pEGFP C2 (addgene: Plasmid #62964, MA, USA). Assembly cloning was performed using pcDNA3-EGFP (addgene: Plasmid #13031, MA, USA) vector inserting BRET from pRetroX-Tight-MCS_PGK-GpNLuc (addgene: Plasmid #70185, MA, USA). Vector was amplified with the primer shown (SI Appendix Table T1. I. a, b of vector) and insert was amplified with primers shown (SI Appendix Table T1. I. a, b inserts) resulting in pcDNA-BRET. The plasmid pcDNA-BRET was amplified with the primers shown (SI Appendix Table T1. I. c, d of vector) and insert CyPET-Arf6 (addgene: Plasmid #18840, MA, USA) was amplified with the primers (SI Appendix Table T1. I. c, d of insert) resulting in SG2. SDM method was exploited to design BMKb9 (MysP-eGFP) by adding the MysP construct upstream of the eGFP cassette in the pcDNA3.1+ vector (addgene: Plasmid #13031, MA, USA) using forward and reverse primer shown (SI Appendix Table T1. II. c). The molecular cloning was validated through Sanger sequencing. Upon publication the plasmids will be made available to other researchers through Addgene and can also be directly shared by the authors upon request.

### 4. Cell Transfection

Transfections were conducted utilizing the chemical transfection reagent Polyethylenimine (PEI) "Max" (catalogue #24765-1, Polysciences, Inc., PA, USA) prepared at a concentration of 1 mg/mL. A 1:3 ratio of DNA to PEI was maintained throughout the transfection procedure. Cells were seeded to achieve approximately 30% confluency prior to transfection. The transfection mixture was prepared in Opti-MEM™ (catalogue # 31985062, Gibco, NY, USA) and incubated at 37 °C for 30 minutes before being added dropwise to the cells, followed by overnight incubation. The media was subsequently replaced with complete media to minimize cellular toxicity. Transfection efficiency was evaluated for 36 hours post-transfection using a CytoFLEX S N2-V3-B5-R3 Flow Cytometer (catalogue #B78557, Beckman Coulter, Inc, IN, USA) and CytExpert software (version 2.4.0.28, Beckman Coulter, Inc, IN, USA). Based on fluorescence signal, samples exhibiting below 75% transfection efficiency were excluded from downstream experiments.

### 5. Confocal Microscopy and Membrane association Measurement

Before analysis by confocal microscopy, the 293A cells were cultured on the surface of gelatin-fibronectin coated glass coverslips (VWR, International, PA, USA) in 24 well plates. The coating solution contained 0.1 % gelatin (Merck Group, MA, USA) and 1 mg/mL fibronectin (Invitrogen, MA, USA). After overnight incubation with coating solution at 37°C, it was washed once with DMEM and 293A cells were seeded to achieve 30% confluency during the transfection. Following 36 hours of incubation, cells were fixed with 4% paraformaldehyde (PFA) (VWR, International, PA, USA) in 0.01 M 1X PBS for 20 minutes at room temperature, followed by extensive washing in 1X PBS. Samples were mounted in ProLong Diamond with 4’,6-diamidino-2-phenylindole (DAPI) (Invitrogen, MA, USA) and examined using a Leica SP8 Lightning confocal microscope. Image analysis and processing was performed using Leica LASX software.

For membrane association analysis, we followed a flowchart (SI Appendix Fig. 8). Membrane and nuclear contours were delineated using ImageJ, and eGFP signal segmentation was conducted with CellProfiler. A custom Python script was developed to determine the proximity of eGFP signals to the membrane and nucleus of individual cells (script available on request). We analyzed the subcellular localization of each marker in cells expressing eGFP-marker fusion constructs generated in-house. We determined the membrane association for each marker by comparing the relative distance to nucleus or membrane, presenting representative images of fluorescence (FL), brightfield (BF), and segmented confocal micrographs of individual cells (Fig. 5A). For this analysis, we utilized the mean of four optimal confocal micrographs exhibiting normal cell morphology. The formula employed to determine the Membrane association percentage was as follows: 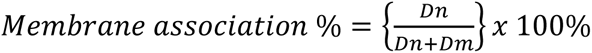

The mean Membrane association % of 4 confocal micrographs of all markers were plotted.

### 6. Isolation of EVs and NVEPs

HEK293 cells (5E+06) were incubated for 24 hours in serum-free media per 150 mm plate having total volume of 20 mL. EVs and NVEPs were isolated with modifications to a previously described method (9, 50). Briefly, cells were removed by centrifugation at 500 ×g for 10 minutes at room temperature, followed by gravity filtration of the supernatant through a 5 µm filter (Merck Group, MA, USA), which was centrifuged (Hermle Z326 K Refrigerated Centrifuge, Hermle AG, Gosheim, Germany) at 2,000 ×g for 20 minutes at 4°C. The supernatant obtained was filtered using 0.8 µm filter (Merck Group, MA, USA) was centrifuged (Hermle Z326 K Refrigerated Centrifuge, Hermle AG, Gosheim, Germany) at 14000 xg for 40 min at 4°C. The pellet obtained was washed with the same centrifugation parameter resulting in 14k-lEV. The supernatant obtained was filtered using 0.2 µm filter (Merck Group, MA, USA) was centrifuged (Optima XPN-100 Ultracentrifuge, B10048, Beckman Coulter, Inc., CA, USA) at 100,000 ×g for 2.5 hours at 4°C. The pellet obtained was washed with the same centrifugation parameter resulting in 100k-sEV and its supernatant was centrifuged (Optima XPN-100 Ultracentrifuge, B10048, Beckman Coulter, Inc., CA, USA) at 167,000 ×g for 16 hours at 4°C and pellet was washed using same centrifugation parameter resulting in 167k-NVEP.

### 7. TEM of EVs and NVEPs

For TEM imaging, we applied the previously described EV isolation method with minor modifications (51). Previously centrifugated concentrated 14k-lEVs, 100k-sEV and 167k-NVEPs pellets were resuspended in 0.1 filtrated 10 µL 1× PBS instead of 50 µL final volume. Samples were fixed in 2% paraformaldehyde (Paraformaldehyde, powder, 95%, 158127, Sigma Aldrich, St. Louis, MO, USA) in 1× PBS for 30 min at 4 °C. Then, they were incubated for 15 min on glow-discharged at 7.2 V for 60 s, carbon-coated 100-mesh copper grids using a Bal-Tec MED 020 Coating System. After washing with 1× PBS and fixating with glutaraldehyde 1% (Sigma-Aldrich, St. Louis, MO, USA) for 5 min at 4 °C, grids were washed again and dried with filter paper. The grids were treated with 2% aqueous uranyl acetate (Sigma-Aldrich, St. Louis, MO, USA) for 2 min to generate negative staining. Samples were washed and dried, then analyzed with FEI Tecnai G2 Spirit TEM (Thermo Fisher Scientific, MA, USA) equipped with a Morada digital camera (Olympus Soft Image Solutions GmbH, Muenster, Germany). All TEM procedures were undertaken at the University of Valencia’s Electron Microscopy facility.

### 8. Size, Concentration, and Zeta Potential Determination of EVs

Size distributions and concentrations of 14k-lEV and 100k-sEV were assessed using nanoparticle tracking analysis (NTA) on a ZetaView PMX120 instrument (Particle Metrix, Germany). Samples were diluted between 1000-fold to 20000-fold in 1X PBS to achieve optimal concentration (50–200 particles/frame) and analyzed across 11 positions with each position was recorded using two sequential measurement cycles for averaging. The following camera settings were used: shutter speed of 100, sensitivity of 70 (for 14k-lEV) and 80 (for 100k-sEV), and a frame rate of 30 frames per second. The zeta potential of 14k-lEV and 100k-sEV was measured three times at 25°C with the following settings: sensitivity of 85, shutter value of 70, and a frame rate of 30 frames per second. ZetaView software (S/N: 19-459 ZetaView Particle Tracking Analyzer PMX-120 instrument, Munich, Germany, employing the ZetaView Analysis 8.05.12 SP2 software) was employed for data collection and analysis.

### 9. Bead-Based Multiplex Flow Cytometry Assay

Bead-based multiplex analysis was conducted using the MACSPlex EV Kit IO human (catalogue # 130-122-209, Miltenyi Biotec, Germany). Assays were performed as described earlier (52, 53). EV and NVEP samples having 4 µg of total protein content measured by micro-BCA kit (catalogue # A55864, Thermo Fisher Scientific, MA, USA), diluted with MACSPlex buffer to a total volume of 120 μl and incubated with 15 μl of MACSPlex exosome capture beads overnight in wells of a pre-wet and drained MACSPlex 96-well 0.22 μm filter plate on an orbital shaker at 450 rpm at room temperature and washed with 200 μl MACSPlex buffer. For counterstaining of captured EVs and NVEPs, a mixture of APC-conjugated anti-CD9, anti-CD63 and anti-CD81 detection antibodies (supplied in the MACSPlex kit, 5 μl each) was added to each well in a total volume of 135 μl, and the plates were incubated on an orbital shaker at 450 rpm for 1 h at room temperature. Next, the samples were washed twice, resuspended in MACSPlex buffer and analysed by flow cytometry with a CytoFLEX S N2-V3-B5-R3 Flow Cytometer (catalogue # B78557, Beckman Coulter, Inc, IN, USA) and CytExpert software (version 2.4.0.28, Beckman Coulter, Inc, IN, USA).

### 10. Single-particle interferometric reflectance imaging sensor (SP-IRIS)

The Leprechaun Exosome Human Tetraspanin capture Kit (catalogue # 251-1044, Unchained Labs, CA, USA) was utilized to capture EVs and NVEPs positive for CD81, CD9, and CD63, following previously established protocols(54). In brief, approximately 1E+06 particles in 50 μL were applied to background-scanned chips and left to incubate for 1 h at room temperature. The Leprechaun instrument scanned the chips using both interferometric microscopy and fluorescence channels corresponding to CD81, CD9, and CD63 signals. Leprechaun Analysis v2.1.1. software was employed to analyze the acquired images for particle concentration and size. 100k-sEV and 167k-NVEP were categorized into seven subpopulations based on their tetraspanin expression: CD9+, CD81+, CD63+, CD9+CD81+, CD9+CD63+, CD63+CD81+, CD9+CD81+CD63+.

### 11. Protein and Lipid Content Determination

The total protein concentration of EV and NVEP preparations was quantified using the Micro BCA Protein Assay kit (catalogue # A55864, Thermo Fisher Scientific, MA, USA) according to the manufacturer’s instructions as previously described (50). EV and NVEP samples were diluted 20-fold to 250-fold and lysed with 0.5% Triton X100 (VWR International, PA, USA). Absorbance was measured at 540 nm on a 96-well plate reader (352 Multiskan MS LabSystems Microplate Reader, 74165-5, Labsystem, Vantaa, Finland). Lipid content of EVs and NVEPs was measured using the sulfo-phospho-vanillin assay as previously described (50, 55). Briefly, EV samples were treated with sulfuric acid, incubated at 90°C, and subsequently reacted with phospho-vanillin reagent (ReagentPlus®, 99%, 79617, Merck, NJ, USA). The resulting colorimetric reaction was measured at 540 nm on a 96-well plate reader (352 Multiskan MS LabSystems Microplate Reader, 74165-5, Labsystem, Vantaa, Finland).

### 12. Raman Measurement

The Raman spectroscopy of EVs and NVEPs was performed by Dr. Jacopo Zini (Timegate Instruments, Oulu, Finland). In brief, the Timegate PicoRaman M3 (Timegate Instruments, Oulu, Finland) with a 100 ps 532 nm laser and complementary metal oxide single-photon avalanche diode was connected to an ÄKTA Pure 25 chromatography system via ProbeProMini (Timegate Instruments, Oulu, Finland) and SCHOTT ViewCell (Schott, Mainz, Germany). Raman spectra were measured continuously throughout the chromatography run with a 20-second acquisition time, 88 mW as the laser power, and a spot size of 100 µm.

### 13. Lipidomics

For quantitative lipidomics, internal standards were added prior to lipid extraction. An amount of 100 µg protein measured by micro-BCA assay was subjected to lipid extraction from HEK293 cells, 100k-sEVs, and 167k-NVEPs were subjected to lipid extraction according to the protocol by Bligh and Dyer (56). The analysis of lipids was performed by direct flow injection analysis (FIA) using a triple quadrupole mass spectrometer (FIA-MS/MS) and a high-resolution hybrid quadrupole-Orbitrap mass spectrometer (FIA-FTMS). FIA-MS/MS was performed in positive ion mode using the analytical setup and strategy described previously (57). A fragment ion of m/z 184 was used for lysophosphatidylcholines (LPC) (58) The following neutral losses were applied: Phosphatidylethanolamine (PE) and lysophosphatidylethanolamine (LPE) 141, phosphatidylserine (PS) 185, phosphatidylglycerol (PG) 189, phosphatidic acid (PA) 115 and phosphatidylinositol (PI) 277 (59). Sphingosine based ceramides (Cer) and hexosylceramides (Hex-Cer) were analyzed using a fragment ion of m/z 264 (60). PE-based plasmalogens (PE P) were analyzed according to the principles described by Zemski-Berry (61). Glycerophospholipid species annotation assumed of even numbered carbon chains only. A detailed description of the FIA-FTMS method was published recently(62). Triglycerides (TG), diglycerides (DG) and cholesterol esters (CE) were recorded in positive ion mode m/z 500-1000 as [M+NH4]+ at a target resolution of 140,000 (at 200 m/z). CE species were corrected for their species-specific response (63). Phosphatidylcholines (PC), PC ether (PC O) and sphingomyelins (SM) were analyzed in negative ion mode m/z 520-960 as [M+HCOO]- at the same resolution setting. Analysis of free cholesterol (FC) was performed by multiplexed acquisition (MSX) of the [M+NH4]+ of FC and the deuterated internal standard (FC[D7]) (63).

LION-PCA heatmap analysis was performed. LION-PCA processes lipidomics datasets by scaling sum-normalized samples as z-scores (mean = 0, standard deviation = 1). PCA is then conducted, and loadings from a selected number of principal components (PrC) are used to rank lipid features. LION term enrichment is evaluated for all selected PrCs using the ranking mode with two-sided KS-tests, as previously described (29). Significantly enriched LION-terms in any of the selected PrCs are identified. The associated lipid feature z-scores for these LION-terms are averaged per sample and visualized in a heatmap using the R package ‘pheatmap’ v1.0.12 (28). Then we compared between 100k-sEV and 167k-NVEP by unpaired two tailed t-test and volcano plot to see enrichment of lipid species between them. Lipidomic enrichment analysis was conducted using LipidSig 2.0, a web-based bioinformatics platform for comprehensive lipid set enrichment and ontology analysis. Following untargeted lipidomic profiling, significantly regulated lipids were uploaded to LipidSig 2.0 for pathway and structural annotation. Enrichment was performed using default settings with lipid ontology categories based on class, subclass, saturation, and chain length.

### 14. High-Resolution Flow Cytometry Measurement

All EV samples were analyzed using the Apogee A60 MP (SN 0130, Apogee Flow Systems, Northwood, UK) running Histogram Software v255.0.0.292 and FCM Control v 3.70. Prior to sample analysis, the system was cleaned using 1% bleach; sheath and diluent were checked to ensure machine and diluent cleanliness (event rate <200 events/second). The Apogee platform was calibrated using a bead mixture (ApoCal 1524, Lot CAL0134, Apogee Flow Systems, Northwood, UK); monitoring beads were also analyzed (Apogee Mix 1527, Lot CAL0139, expiration 20-SEP-2021, Apogee Flow Systems, Northwood, UK) as part of daily protocols. Samples were analyzed for 60 seconds at a 3.01 µL/min flow rate, recording approximately 80,000 events per run. The analysis was performed using FlowJo 10.9.0.

### 15. Super-Resolution Microscopy Measurement

Samples were diluted tenfold in 1X PBS, placed onto cover glasses, and incubated overnight at 4°C in humidity chambers. Annexin V Alexa Fluor 647 conjugate (Invitrogen, MA, USA, R37175) was added to the samples on coverslips (VWR International, PA, USA) and incubated for 15 minutes at room temperature, protected from light. For fixation, 4% PFA-PBS was used for 5 minutes at room temperature, followed by three washing steps of 5 minutes each with PBS. Antigen retrieval was carried out using 0.001% Triton X100 (T9284-500ML; Merck Group, MA, USA) in PBS for 5 minutes at room temperature, followed by 3 washing steps (3 x 5 minutes). Non-specific binding sites were blocked by 5% bovine serum albumin (BSA) in PBS at 37°C for 35 minutes. All primary antibodies: anti-GFP antibody (Catalog # A11122, Invitrogen, MA USA), anti-HSP90ß antibody (Catalog # 37-9400, Invitrogen, MA USA) were diluted at a 1:200 ratio in 10% BSA-PBS and applied to the samples for 1-hour incubation at room temperature, followed by 3 washing steps. Goat anti-Rabbit Alexa Fluor 568 (Catalog # A11011, Invitrogen, MA, USA) was applied at a 1:400 dilution at room temperature for 1 hour, protected from light. After another washing step with sterile PBS (5 minutes), the cover glasses were placed on cavity slides (BR475505-50EA, Merck Group, MA, USA) filled with blinking buffer and sealed with two-component adhesive. Blinking buffer contained 100 U glucose oxidase (G2133-50KU, Merck Group, MA, USA), 2000 U CAT (C100, Merck Group, MA, USA), 55.56 mM glucose (G8270, Merck Group, MA, USA), and 100 mM cysteamine hydrochloride (M6500, Merck Group, MA, USA) in 1 mL final volume with sterile PBS. The dSTORM imaging was performed using a Nanoimager S (Oxford Nanoimaging ONI Ltd, Oxford, UK) instrument and evaluated by a built-in application of the corresponding quantification software CODI, called EV profiling, which identifies subpopulations with single EV particle using a customized clustering workflow shown in SI Appendix Fig. 9. Cluster sizes were processed accounting for the size of primary antibodies with an average length of about 14 nm.

### 16. Functional Uptake Assay

HEK293T cells were transfected with plasmids encoding eGFP-tagged versions of Arf6, CD63, LC3b, MysP, and Rab7a, following the protocol detailed in the Cell Transfection section. The 167k-NVEPs were subsequently isolated from transfected cells as described in the Isolation of EVs and NVEPs section. Total protein content of each 167k-NVEP preparation was quantified using the micro-BCA assay (Micro BCA™ Protein Assay Kit, 500 mL per kit, Thermo Fisher Scientific, MA, USA). For the functional uptake assay, HEK293T cells seeded in 96 well plates were treated with increasing amounts of 167k-NVEPs—specifically, 1 μg, 5 μg, 20 μg, 100 μg, and 500 μg total protein per well. Cells were incubated with NVEPs for 6 hours under standard culture conditions. Following incubation, cells were harvested and analyzed by flow cytometry to quantify the percentage of eGFP-positive cells (eGFP%). Measurements were performed using the CytoFLEX S N2-V3-B5-R3 Flow Cytometer (B78557, Beckman Coulter, Brea, USA) and CytExpert software (version 2.4.0.28, Beckman Coulter, Inc., Brea, USA).

### 17. Statistical Analysis

In this study, unpaired two-tailed t-tests and Mann-Whitney tests were conducted for comparison between two groups. For comparisons involving more than two groups, the dataset was initially assessed for normality and lognormality. Based on the results, either an ordinary one-way ANOVA with Tukey’s multiple comparisons test or the Kruskal-Wallis test was employed. For more than 3 groups we employed ordinary two-way ANOVA The analysis was performed utilizing GraphPad Prism 10 (GraphPad Software, La Jolla, USA). Deconvolution of the Raman spectroscopy was performed using OriginPro 2019 (OriginLab, Northampton, MA, USA), and other specialized software as appropriate. PCA was done using the Statistics and Machine Learning Toolbox™ of MATLAB version: 9.7.0.1190202 R2019b (MathWorks). PCA of both lipidomic and Raman data was done using singular value decomposition algorithm due to the number of variables exceeding the number of observations. Based on the Keiser method, the lipidomic data has only two meaningful components and the Raman data three, however, for the sake of easily interpretable visualization, the same parameters for both analyses were implemented – that is, the number of requested components was set to 3. Variables were centered prior to transforming the data into a reduced subspace. The code is available upon request.

## Acknowledgments

We thank our Semmelweis University colleagues; Dr. Angéla Takács for sharing the EBC1 cell line, and Dr. Péter Várnai for sharing the 293A cell line. We also extend our sincere gratitude to Mario Soriano (Centro Investigácion Príncipe Felipe, Valencia, Spain) for his assistance with TEM, and Dr. Jacopo Zini (Timegate Instruments, Oulu, Finland) for support with Raman spectroscopy.

The project has received funding from the EU’s Horizon 2020 Research and Innovation Programme under Grant agreement No. 739593, OTKA FK 147023, EXCELLENCE 151417, Advanced Grant 150767, 2019-2.1.7-ERA-NET-2021-00015, and NVKP_16-1-2016-0004 grants of the Hungarian National Research, Development and Innovation Office (NKFIH), the Higher Education Excellence Program (FIKP) and the Therapeutic Thematic Programme TKP2021-EGA-23, RRF-2.3.121-2022-00003 (National Cardiovascular Laboratory Program), VEKOP-2.3.2-162016-00002, VEKOP-2.3.3-15-2017-00016, EKÖP-2024-237 and Stipendium Hungaricum Scholarship 2021.

