## Supplementary material for "Cholesteryl esters and high protein-to-lipid ratios distinguish Non-Vesicular Extracellular Particles from Extracellular Vesicles": SI Appendix

Affiliations:
1. HCEMM-SU Extracellular Vesicle Research Group, Budapest, 1089, Hungary
2. Institute of Genetics, Cell- and Immunobiology, Semmelweis University, Budapest, 1089, Hungary
3. Division of Medical Oncology, Modena University Hospital, Modena, 41124, Italy
4. First Department of Internal Medicine, University of Szeged, Szeged, 6725, Hungary
5. Institute of Behavioural Sciences, Semmelweis University, Budapest, 1089, Hungary
6. Department of Pathology and Experimental Cancer Research, Semmelweis University, Budapest, 1085 Hungary
7. Institute of Clinical Chemistry and Laboratory Medicine, Regensburg University Hospital, Regensburg, 93053, Germany
8. HUN-REN-SU Translational Extracellular Vesicle Research Group, Budapest, 1089, Hungary
* Corresponding author

#### 8. Lipidomics

Lipid extraction from cells, 100k-sEVs, and 167k-NVEPs (equivalent to 100 μg of total protein), along with subsequent processing and quantification via lipid mass spectrometry for cholesterol, glycerophospholipids, and primary sphingolipids, was conducted as previously described (6, 7). A hybrid quadrupole-Orbitrap mass spectrometer utilizing Fourier-Transform mass spectrometry and a triple quadrupole mass spectrometer employing tandem mass spectrometry were utilized to measure lipid species as described earlier (8, 9). The analysis encompassed five broad lipid categories: Fatty Acyls, Glycerolipids, Glycerophospholipids, Sphingolipids, and Sterol Lipids. Custom-programmed computer macros (Excel, Microsoft Corp., Redmond, WA) were employed for deisotoping and data analysis (10). Lipid species were labeled according to the recently proposed shorthand notation for lipid structures derived from mass spectrometry (11). Additional LION-PCA heatmap analysis was performed. LION-PCA processes lipidomics datasets by scaling sum-normalized samples as z-scores (mean = 0, standard deviation = 1). PCA is then conducted, and loadings from a selected number of principal components (PrC) are used to rank lipid features. LION term enrichment is evaluated for all selected PrCs using the ranking mode with two-sided KS-tests, as previously described (12). Significantly enriched LION-terms in any of the selected PrCs are identified. The associated lipid feature z-scores for these LION-terms are averaged per sample and visualized in a heatmap using the R package 'pheatmap' v1.0.12 (13). Then we compared between 100k-sEV and 167k-NVEP by unpaired two tailed t-test and volcano plot to see enrichment of lipid species between them. Lipidomic enrichment analysis was conducted using LipidSig 2.0, a web-based bioinformatics platform for comprehensive lipid set enrichment and ontology analysis. Following untargeted lipidomic profiling, significantly regulated lipids were uploaded to LipidSig 2.0 for pathway and structural annotation. Enrichment was performed using default settings with lipid ontology categories based on class, subclass, saturation, and chain length.

#### 12. Molecular Cloning

In this study, we have used 2 recombinant plasmids, which was developed in-house. They are Myc_EGFP_Arf6_HA (SG2) and MysP-eGFP (SG5). We have also used pEGFP-LC3 (addgene: Plasmid #24920, MA, USA), GFP-rab7 WT (addgene: Plasmid #12605, MA, USA), and CD63-pEGFP C2 (addgene: Plasmid #62964, MA, USA). Assembly cloning was performed using pcDNA3-EGFP (addgene: Plasmid #13031, MA, USA) vector inserting BRET from pRetroX-Tight-MCS_PGK-GpNLuc (addgene: Plasmid #70185, MA, USA). Vector was amplified with the primer shown (Extended Table T1. I. a, b of vector) and insert was amplified with primers shown (Extended Table T1. I. a, b inserts) resulting in pcDNA-BRET. The plasmid pcDNA-BRET was amplified with the primers shown (Extended Table T1. I. c, d of vector) and insert CyPET-Arf6 (addgene: Plasmid #18840, MA, USA) was amplified with the primers (Extended Table T1. I. c, d of insert) resulting in SGb42. SDM method was exploited to design BMKb9 (MysP-eGFP) by adding the MysP construct upstream of the eGFP cassette in the pcDNA3.1+ vector (addgene: Plasmid #13031, MA, USA) using forward and reverse primer shown (Extended Table T1. II. c). The molecular cloning was validated through Sanger sequencing.

#### 13. Membrane association analysis

For membrane association analysis, we followed a flowchart (SI Appendix Fig. 8). Membrane and nuclear contours were delineated using ImageJ, and eGFP signal segmentation was conducted with CellProfiler. A custom Python script was developed to determine the proximity of eGFP signals to the membrane and nucleus of individual cells (script can be found in S8.docx). We analyzed the subcellular localization of each marker in cells expressing eGFP-marker fusion constructs generated in-house. We determined the membrane association for each marker by comparing the relative distance to nucleus or membrane, presenting representative images of fluorescence (FL), brightfield (BF), and segmented confocal micrographs of individual cells (Fig. 5a). For this analysis, we utilized the mean of four optimal confocal micrographs exhibiting normal cell morphology. The formula employed to determine the membrane proximity percentage was as follows: $Membrane\ proximity\ \% = \left\{\frac{Dn}{Dn+Dm}\right\} x\ 100\%$

The mean membrane proximity % of 4 confocal micrographs of all markers were plotted.

**Supplementary Figure 1.**

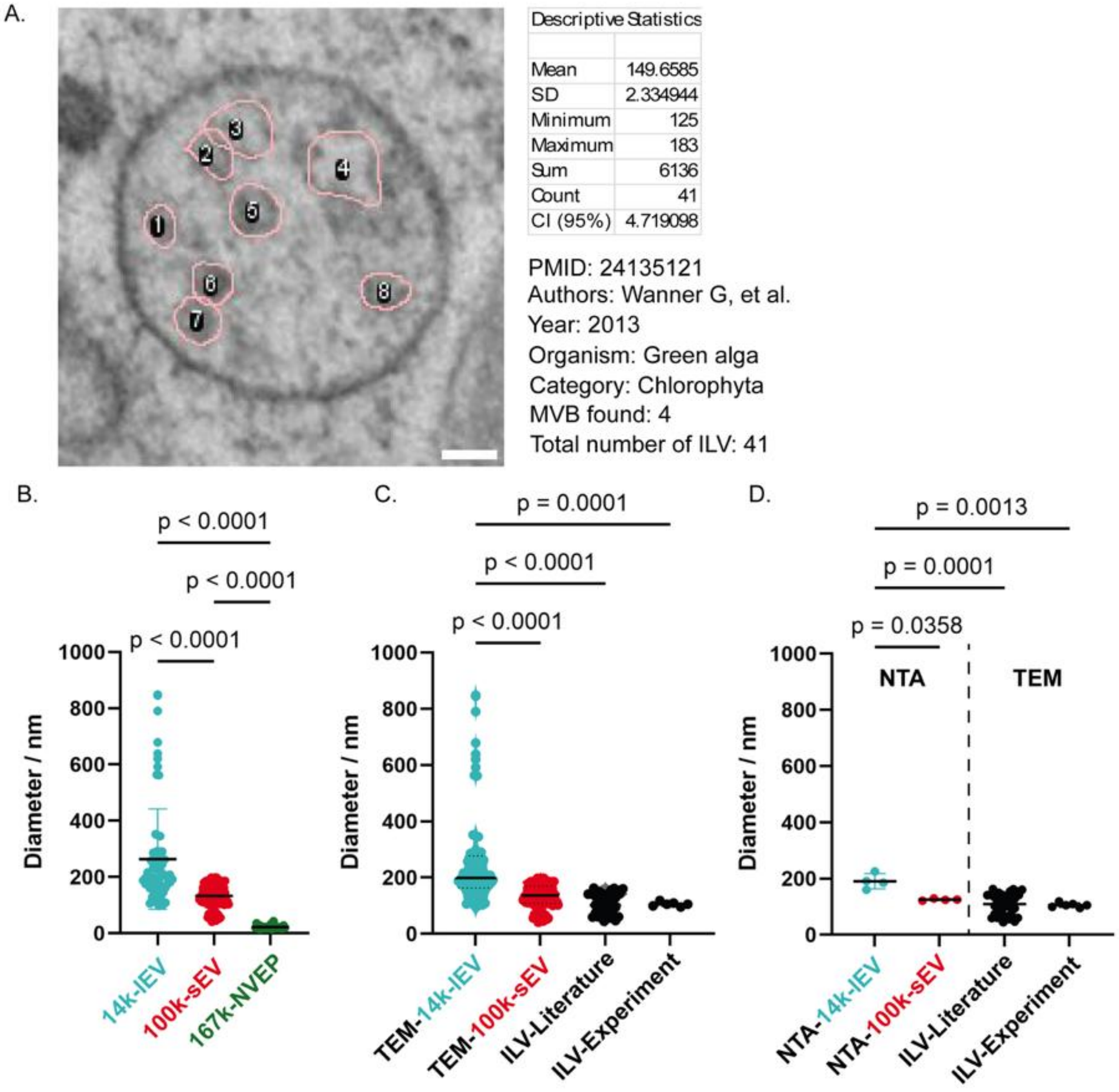


**Supplementary Figure 1. Image analysis and TEM-based characterization of ILV and EV subpopulation sizes.**

(A) To estimate intraluminal vesicle (ILV) diameters, image analysis was performed on transmission electron microscopy (TEM) micrographs from 39 studies meeting meta-analysis inclusion criteria. Using ImageJ, the Feret diameter (nm) was measured by outlining the ILV boundaries. For each study, up to 152 ILVs were quantified, and descriptive statistics were calculated. A representative ImageJ-based analysis of a single multivesicular body (MVB) is shown, with associated statistical parameters. (B) TEM-based size measurements of extracellular vesicle (EV) subtypes isolated from HEK293 cells revealed significant differences: 14k-lEV (263.90 ± 178.10 nm), 100k-sEV (133.20 ± 42.04 nm), and 167k-EP (21.74 ± 6.37 nm) (mean ± standard deviation; $p<0.0001$, ordinary one-way ANOVA; $n\geq31$ per group). (C) Comparison of TEM-derived diameters showed that 14k-lEVs were significantly larger than both 100k-sEVs and ILVs (from both experimental and literature-derived data) ($p \leq 0.0001$, ordinary one-way ANOVA). (D) We compared the sizes of 14k-lEV and 100k-sEV as measured by NTA. NTA analysis showed a significant size difference between 14k-lEV and 100k-sEV ($p=0.0358$, ordinary one-way ANOVA). Furthermore, 14k-lEVs measured by NTA were significantly larger than ILVs measured by TEM from both our experimental dataset ($p=0.0001$, ordinary one-way ANOVA) and literature-derived data ($p=0.0013$, ordinary one-way ANOVA).

**Supplementary Figure 2.**

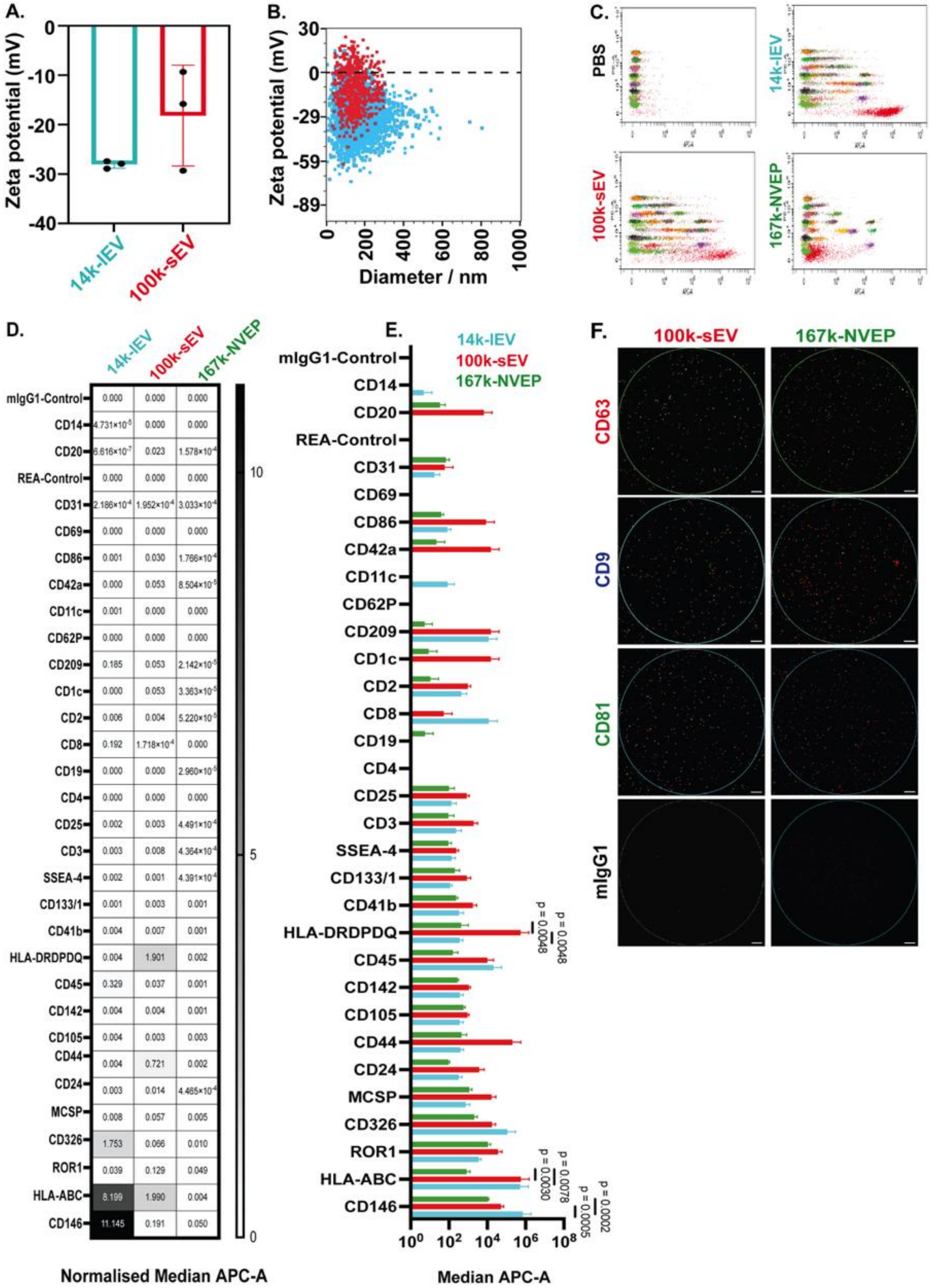

**Supplementary Figure 2 (related to figure 2). Characterization of individual EV subpopulations using multiplex bead-based flow cytometry and SP-IRIS.**

(A) Zeta potential measurements showing -28.07 ± 0.76 mV for 14k-lEV and -18.13 ± 10.20 mV for 100k-sEV (n=3 biological replicates). (B) Scatterplot depicting zeta potential as a function of particle size for individual 14k-lEV (cyan) and 100k-sEV (red) fractions (n=4 biological replicates). (C) Surface marker profiling of EV subpopulations—14k-lEV, 100k-sEV, and 167k-NVEP—was performed using multiplex bead-based flow cytometry (MACSPlex EV Human Kit). A representative gating strategy is shown, with PE-A vs. APC-A plots. Phosphate-buffered saline (PBS) served as a negative control, and gating thresholds were set to exclude background signals arising from the PBS region. (D) To enable inter-marker comparison, median APC-A fluorescence intensities were normalized by calculating the ratio of each marker signal to the average of canonical EV tetraspanins (CD9, CD63, and CD81). Following normalization, no substantial expression differences were observed for most markers across EV subtypes, except for CD146, which was elevated in 14k-lEV, and HLA-ABC, which was enriched in both 14k-lEV and 100k-sEV. (E) Among the 33 markers assessed, low-intensity APC-A signals were generally observed. Notably, CD146 expression was significantly higher in 14k-lEV, and HLA-DRDPDQ was enriched in 100k-sEV ($p < 0.01$, two-way ANOVA; n=3 biological replicates). (F) Single-particle interferometric reflectance imaging sensing (SP-IRIS) validated the presence of CD9, CD63, and CD81 on both 100k-sEV and 167k-NVEP subpopulations. No detectable signal was observed in the mIgG1 isotype control. A representative image from three independent biological replicates is shown. Data are presented as mean ± standard deviation, where applicable.

**Supplementary Figure 3.**

A.

| Raman Shift (cm$^{-1}$) | Functional Group | Vibrations/Assignment |
|---|---|---|
| **701** | cholesteryl ester | Cholesterol ring deformation |
| **760** | phosphodiester | Nucleic acid |
| **783** | aromatic residue | Trp, side chain of protein |
| **850** | C-C | Protein backbone |
| **1006** | aromatic residue | Phe, side chain of protein |
| **1055-1070** | C–C vibrations of lipids | Lipids I |
| **1130** | aromatic residue | Phe, side chain of protein |
| **1210** | C–C twisting | Lipids, triacyl glycerol |
| **1340** | δ C-H (CH2) | Glycosaminoglycans |
| **1441** | CH2/CH3 | Saturated lipids |
| **1556** | Indole ring | Tryptophan |
| **1661** | Amide I | Proteins |
| **1720** | Ester bond | Lipids |

B.

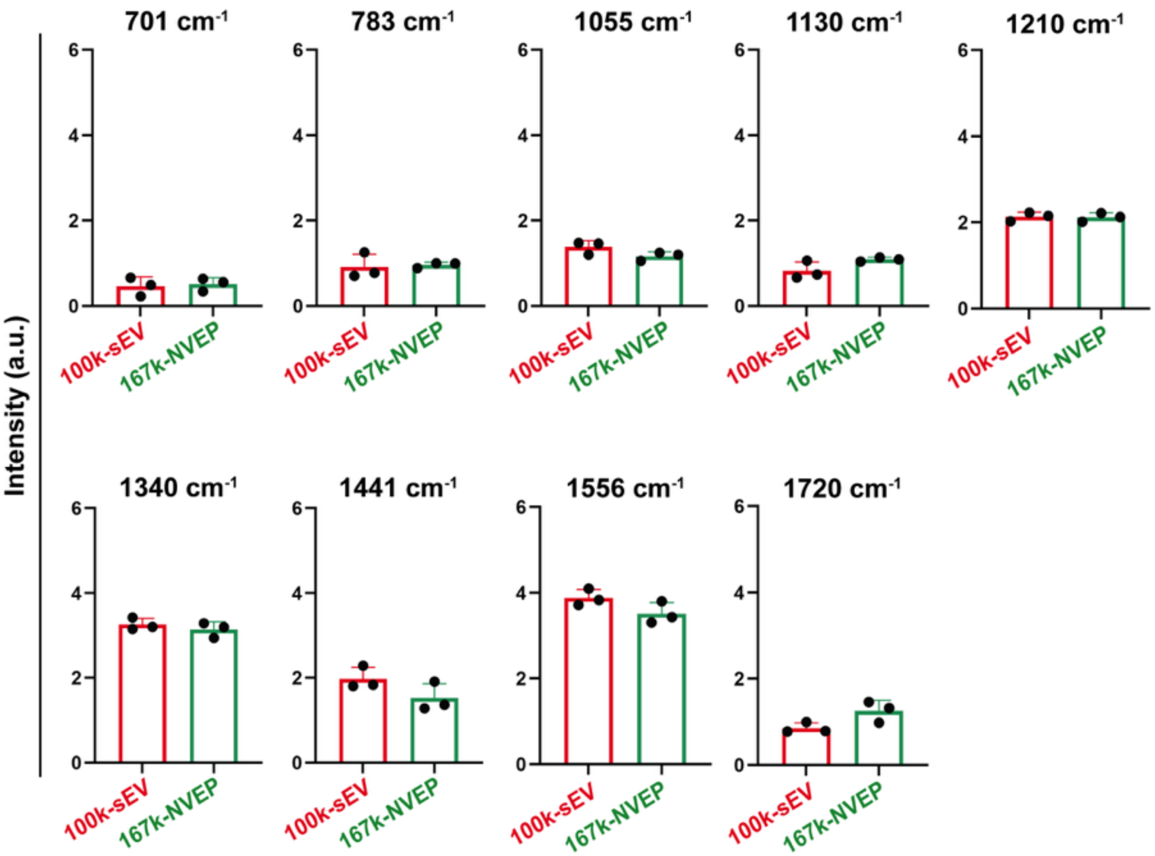

**Supplementary Figure 3 (related to figure 3). Raman spectroscopic profiling of HEK293-derived extracellular vesicles.**

(A) Representative Raman vibrational band assignments for HEK293-derived 100k-sEV and 167k-NVEP were based on established literature (16, 17). Spectral peaks correspond to characteristic molecular components such as lipids, nucleic acids, and proteins. (B) Comparative analysis of spectral fingerprints revealed significant differences between 100k-sEV and 167k-NVEP at several key Raman shifts: 708 $cm^{-1}$ (cholesterol ring deformation), 783 $cm^{-1}$ (nucleic acids), 1055 $cm^{-1}$ (Lipids I), 1130 $cm^{-1}$ (phenylalanine, protein side chain), 1210 $cm^{-1}$ (triacylglycerol), 1340 $cm^{-1}$ (glycosaminoglycans), 1441 $cm^{-1}$ (saturated lipids), 1556 $cm^{-1}$ (tryptophan), and 1720 $cm^{-1}$ (lipids), with $p<0.05$ for all comparisons (unpaired t-test; n=3 biological replicates).

**Supplementary Figure 4.**

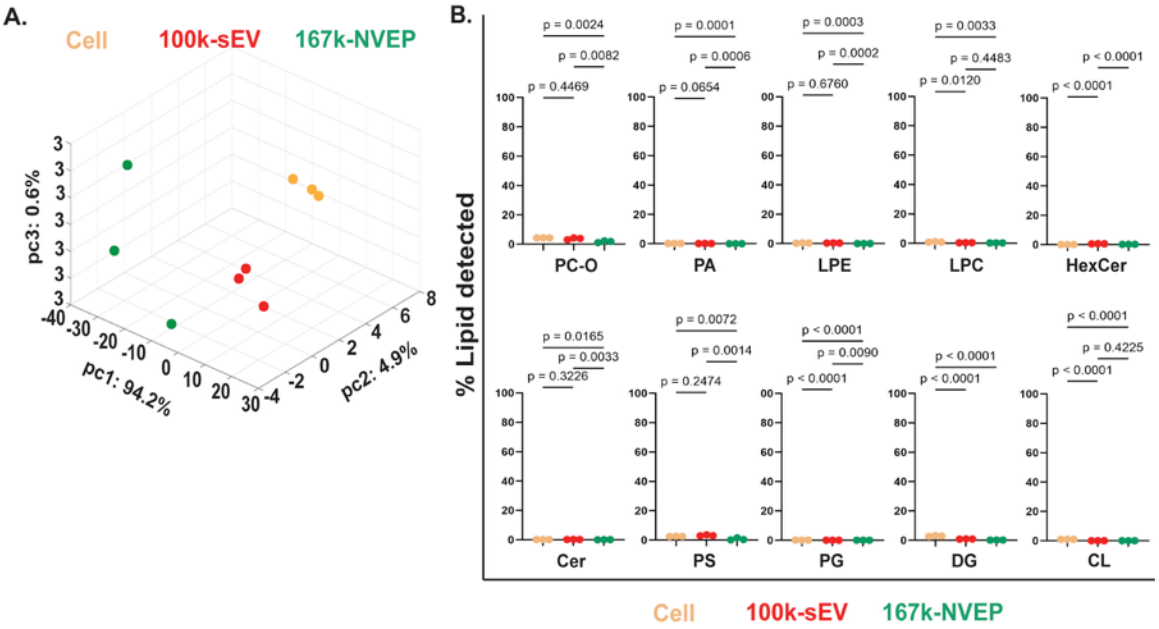


**Supplementary Figure 4 (related to figure 4). Comprehensive lipidomic profiling of cells, 100k-sEV, and 167k-NVEP**

(A) Three-dimensional principal component analysis (PCA) plot illustrating distinct lipid signatures of cells (yellow), 100k-sEV (red), and 167k-NVEP (green). Each point represents an independent biological replicate (n=3). Axes correspond to principal components 1 (94.2%), 2 (4.9%), and 3 (0.6%), capturing 99.7% of total variance. Analysis includes eight EV-associated lipid classes: free cholesterol (FC), sphingomyelin (SM), phosphatidylethanolamine plasmalogen (PE-P), cholesteryl ester (CE), phosphatidylcholine (PC), phosphatidylethanolamine (PE), phosphatidylinositol (PI), and triglycerides (TG). Data were processed using MetaboAnalyst 5.0 with log transformation and Pareto scaling (FDR<0.05). Clustering aligns with MISEV2023 (18) guidelines for distinguishing EV subtypes and non-vesicular particles. (B) Relative abundance of specific lipid species across cells, 100k-sEV, and 167k-NVEP. Lipid species including phosphatidylcholine-ether (PC-O), phosphatidic acid (PA), lysophosphatidylethanolamine (LPE), lysophosphatidylcholine (LPC), hexosylceramide (HexCer), ceramide (Cer), phosphatidylserine (PS), phosphatidylglycerol (PG), diglyceride (DG), and cardiolipin (CL) show lower abundance in cells compared to their secreted 100k-sEV and 167k-NVEP counterparts. Values are normalized peak intensities (log2-transformed) from n=3 biological replicates.

**Supplementary Figure 5.**

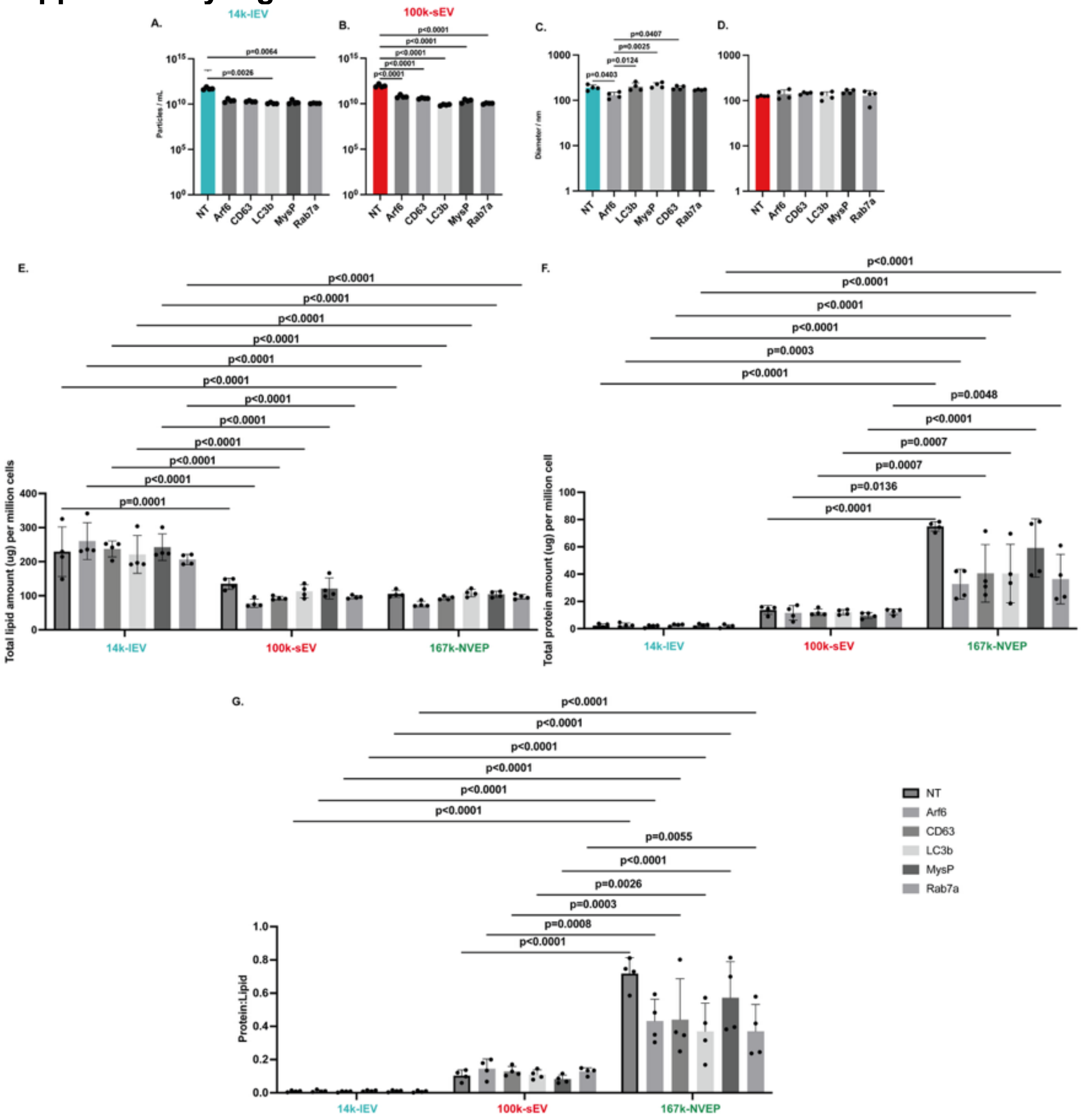


### Supplementary Figure 5. Characterization of Marker-Expressing EVs

(A) LC3b, and Rab7a expressing 14k-lEVs showed significantly lower concentrations compared to their non-transfected counterparts (p=0.0057, 0.0098, and 0.0015, respectively, Ordinary one-way ANOVA). (B) All marker-expressing 100k-sEVs exhibited significantly lower concentrations compared to non-transfected 100k-sEVs (p<0.0001, Ordinary one-way ANOVA). (C) The mean size of all 14k-lEVs was 187.0 ± 34.99 nm. (D) All 100k-sEVs averaged 132.2 ± 13.50 nm, with no significant differences in sizes between any of the marker-expressing variants. (E) Total lipid of 14k-lEV per million Hek293 cells was significantly higher compared to both 100k-sEV and 167k-NVEP (p<0.0001, Two-way ANOVA, n=4). (F) Total protein of 167k-NVEP per million Hek293 cells was significantly higher compared to both 14k-lEV and 100k-sEV (p<0.05, Two-way ANOVA, n=4). (G)

167k-NVEPs showed have significantly high protein to lipid ratio than EV subtypes ($p<0.01$, Two-way ANOVA, $n=4$).

**Supplementary Figure 6**.

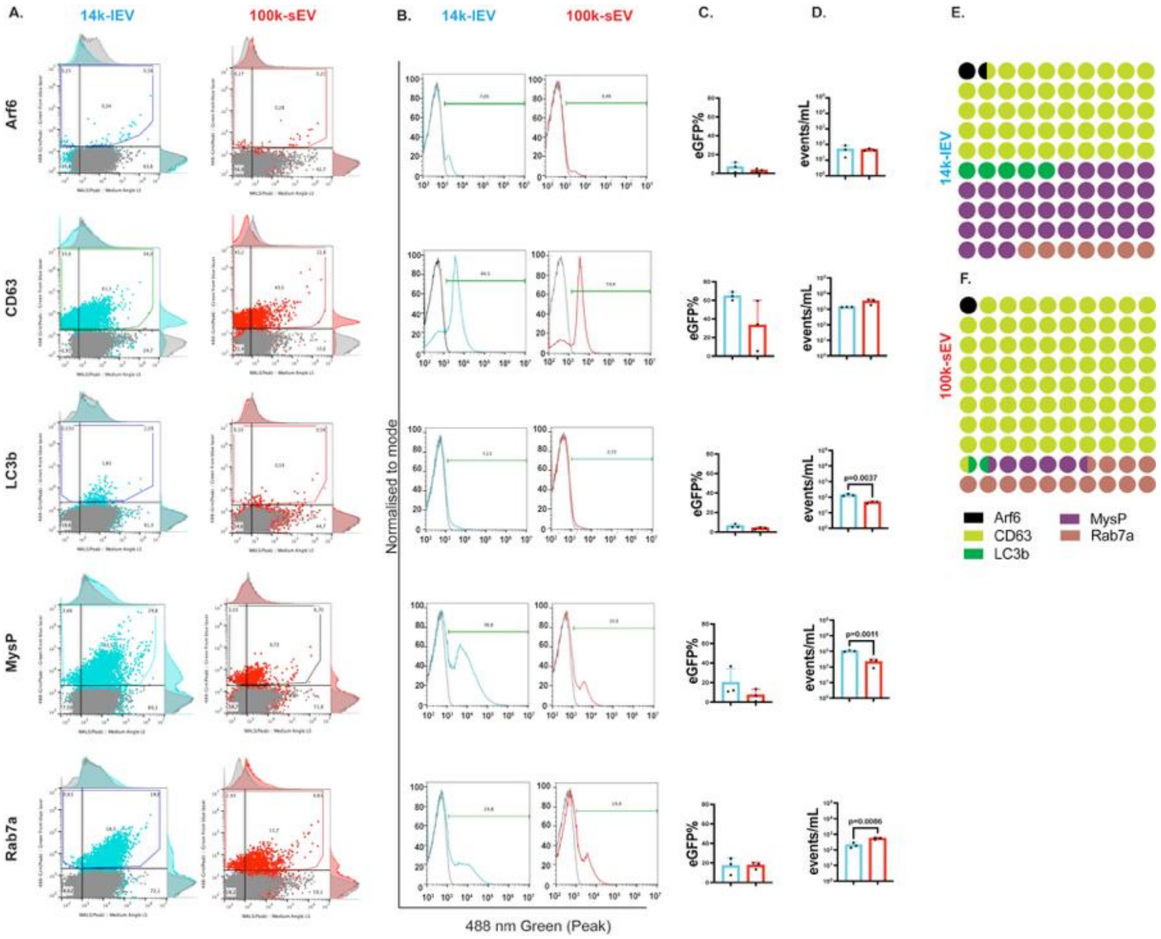


**Supplementary Figure 6 (related to figure 6)**. **High resolution Flow Cytometry Analysis of EVs**

(A) Flow cytometry data were obtained from the Apogee A60 MP (SN 0130, Apogee Flow Systems, Northwood, UK) and analyzed using FlowJo 10.9.0 software. Gating was performed on native 14k-lEV and 100k-sEV samples that did not express any markers or eGFP. Signals from non-transfected 14k-lEV and 100k-sEV were excluded from the gating analysis, and a single representative plot is shown. Data represent n = 3 biological repeats. (B) Representative of high-resolution flow cytometry histograms showing single-vesicle detection of eGFP-tagged markers, including Arf6, CD63, LC3b, MysP, and Rab7a for 14k-lEV (cyan) and 100k-sEV (red) populations. (C) Percentage of eGFP-positive events for each marker in 14k-lEV and 100k-sEV, where similar distribution of data was observed. (D) Absolute counts of eGFP-positive events per mL for each marker, highlighting subtype-specific enrichment: LC3b, MysP in 14k-lEV and Rab7a, in 100k-sEV (unpaired two-tailed t test, $p<0.01$). Matrix dot plots showing mean of events/mL for each marker: Arf6 (black), CD63 (chartreuse), LC3b (green), MysP (purple), and Rab7a (coral) in 14k-lEV (d) and 100k-sEV (E). No marker was exclusive to a single subtype, but CD63 (48.43%) and MysP (38.16%) were more abundant in

14k-lEV, (F) while CD63 (79.28%) and Rab7a (13.47%) were more abundant in 100k-sEV. All data are shown as mean ± standard deviation from three independent experiments.

**Supplementary Figure 7.**

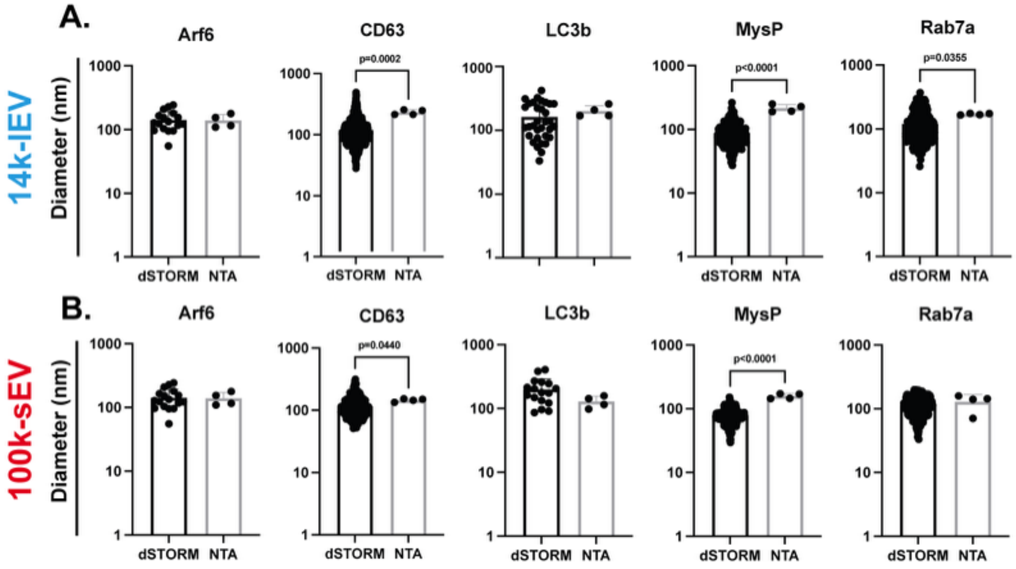


**Supplementary Figure 7. Size Comparison of EVs Using NTA and dSTORM**

(A) Diameter of 100k-sEVs expressing different markers was compared using two orthogonal methods: NTA and dSTORM. The median diameter of 14k-lEVs expressing CD63 (190.70 ± 18.10 nm), MysP (215.80 ± 29.43 nm), and Rab7a (170.50 ± 4.81 nm) obtained by NTA was significantly higher than that measured by dSTORM ($p<0.05$, Mann-Whitney test). (B) A similar trend was observed for the median diameter measured by NTA for 100k-sEVs expressing CD63 (145.20 ± 8.29 nm) and MysP (158.10 ± 14.96 nm), which were significantly higher than the measurements obtained by dSTORM ($p<0.05$, Mann-Whitney test).

**Supplementary Figure 8**.

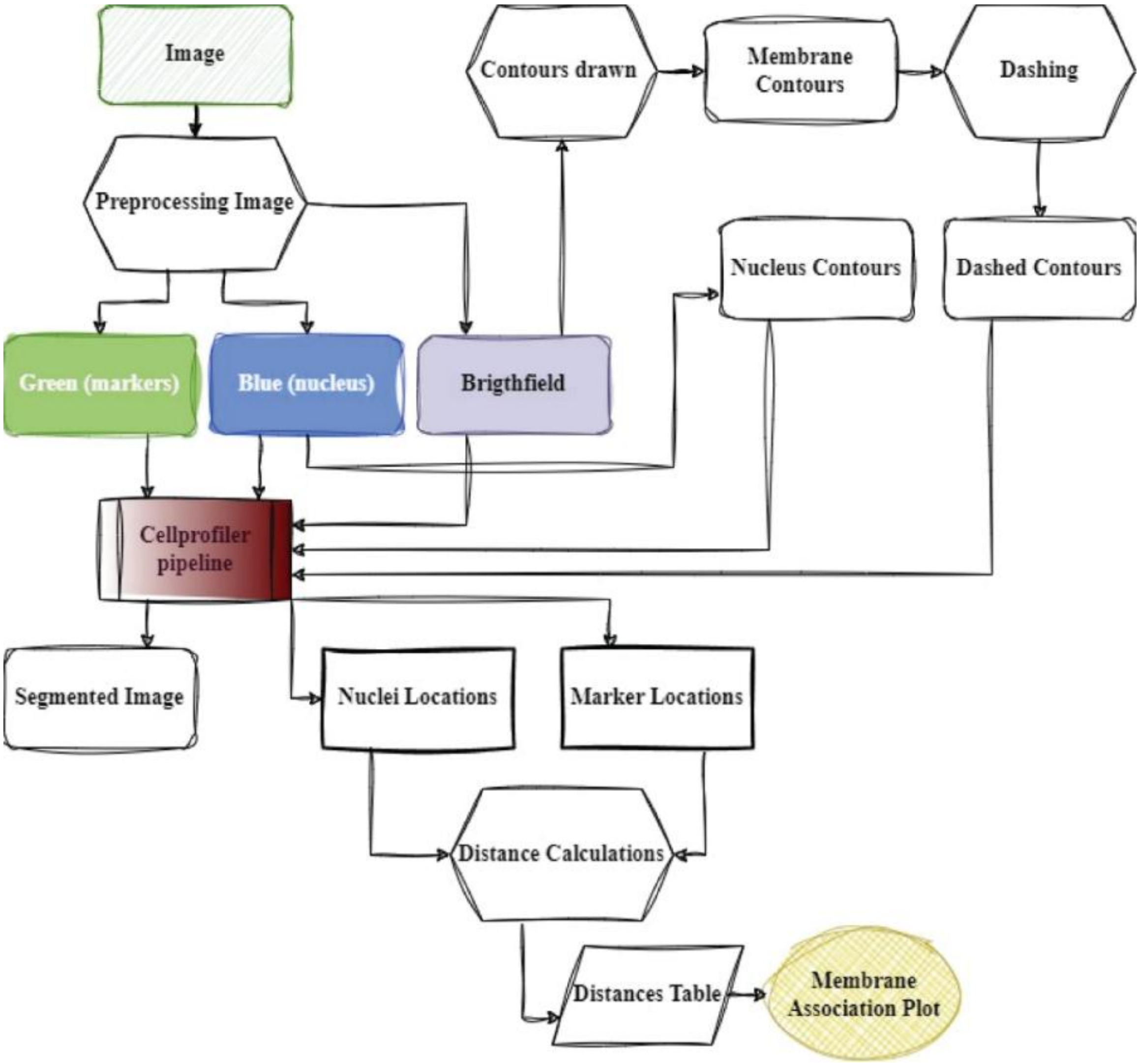


**Supplementary Figure 8 (related to figure 5)**. **Image Processing and Analysis Pipeline**

The entire pipeline (designed in draw.io) is shown, starting from the acquisition of fluorescence and brightfield images of individual 293A cells expressing various markers, such as Arf6, CD63, LC3b, MysP, Rab7a, Syn1, and Tspan14. Images were preprocessed using Python, and contours were drawn from the nucleus and brightfield images to obtain the nucleus and membrane contours, which are represented as dashed lines. These contour images were analyzed in CellProfiler 4.2.8, where the eGFP signal was segmented to determine the nucleus location and marker location (eGFP signal location in the cell). Distances between the nucleus and markers were calculated, generating a distance table using Python, and a membrane association plot was created using GraphPad Prism 10.3.0.

**Supplementary Figure 9**.

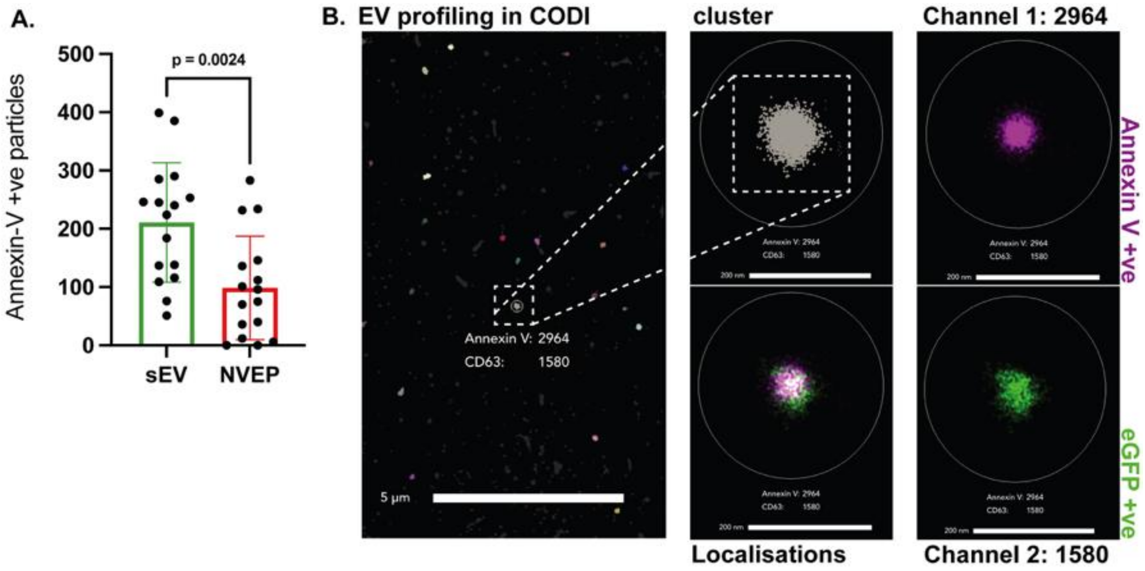


**Supplementary Figure 9 (related to figure 7). dSTORM analysis of the EPs**
(A) Bar plot showing that native 167k-NVEPs has significantly less Annexin-V signal than 100k-sEV ($p<0.0024$, Two-tailed Unpaired t test, $n=16$). (B) Representative dSTORM images of 14k-lEVs expressing CD63, where AnnexinV-positive and eGFP-positive signals are detectable. Each cluster was analyzed using the built-in EV profiling application of the CODI quantification software, which identifies subpopulations of single EV particles through a customized clustering workflow. This workflow first identifies clusters merging the signals of AnnexinV and eGFP. Here, a single 100k-sEV expressing CD63 contains 2964 AnnexinV-positive signals and 1580 eGFP-positive signals. Scale bar 5 μm (left) 200 nm (center and right).

**Supplementary Figure 10.**

.

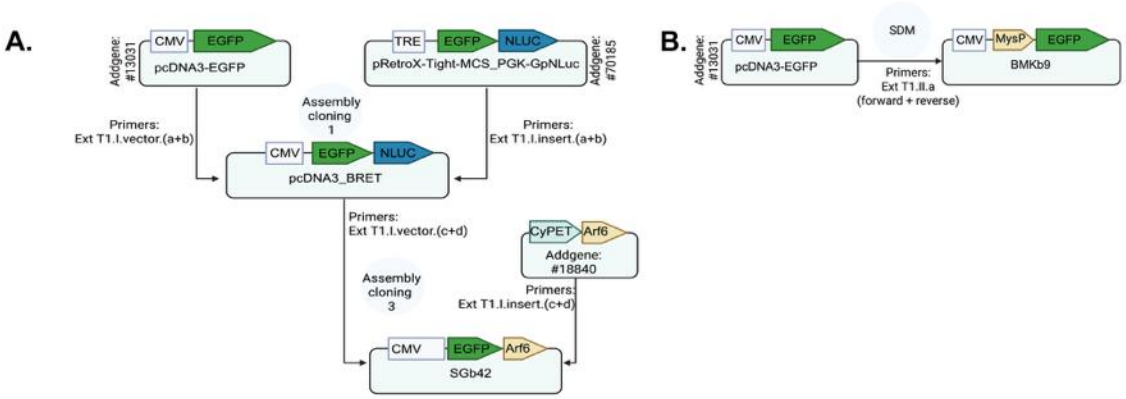


**Supplementary Figure 10. Schematic overview of the molecular cloning strategy**

(A) Assembly cloning was initiated using a construct obtained from Addgene to generate the pcDNA3_BRET vector, which served as the backbone for subsequent cloning steps. This vector was then used to generate constructs SG2 (eGFP_ARF6) for downstream experiments, (B) SDM was further used to introduce a myristoylation and palmitoylation (MysP) cassette upstream of the eGFP coding region named as SG5 (MysP_eGFP). Details of the primers used for each cloning and mutagenesis step, as well as the final constructs, are provided in the Extended Table T1 and T2.

**Supplementary Table T1. Overview of primers utilized for assembly cloning and SDM.**

| I. | Vector (5'-3') | Insert (5'-3') |
|---|---|---|
| a | GCAGCGGCGGCAGCGGCGGCGTGAGCAAGGGCGAGGAG | GGAGGATCAGGAGGCAGTGGTGGCGGTAC |
| b | CCACTGCCTCCTGATCCTCCCGCCAGAATGCGTTCGCAC | GCCGCCGCTGCCGCCGCT |
| c | GGAGGATCAGGAGGCAGTG | ACAAGGATATCTCCGGAGGTGGGAAGGTGCTATCCAAAATC |
| d | ACCTCCGGAGATATCCTTG | CCACTGCCTCCTGATCCTCCCACACTTCCAGCACTCGAG |
| **II.** | **Forward Primer** | **Reverse Primer** |
| a | CAAGAGAAAGGACGTGAGCAAGGGCGAGGAGC | CTGTTGCAGCAGCCCATCTCGAGCGGCCGCCA |

**Supplementary Table T2. Description of plasmids used, including specific single amino acid sequences. Distinct regions are highlighted with various colors for clarity.**

| Plasmid ID | Name and amino acid sequence |
|---|---|
| SG2 | Myc_EGFP_Arf6_HA |
| | EQKLISEEDLGSGGSGGSGGVSKGEELFTGVVPILVELDGDVNGHKFSVSGEGEGDATYG<br>KLTLKFICTTGKLPVPWPTLVTTLTYGVQCFSRYPDHMKQHDFFKSAMPEGYVQERTIFFK<br>DDGNYKTRAEVKFEGDTLVNRIELKGIDFKEDGNILGHKLEYNYNSHNVYIMADKQKNGIK<br>VNFKIRHNIEDGSVQLADHYQQNTPIGDGPVLLPDNHYLSTQSALSKDPNEKRDHMVLLEF<br>VTAAGITLGMDELYKDISGGGKVLSKIFGNKEMRILMLGLDAAGKTTILYKLKLGQSVTTIPT<br>VGFNVETVTYKNVKFNVWDVGGQDKIRPLWRHYYTGTQGLIFVVDCADRDRIDEARQELH<br>RIINDREMRDAIILIFANKQDLPDAMKPHEIQEKLGLTRIGSSAGSVGGSGGSGGGTYPYDV<br>PDYA |
| SG5 | MysP-eGFP |
| | MGCCNSKRKDVSKGEELFTGVVPILVELDGDVNGHKFSVSGEGEGDATYGKLTLKFICTT<br>GKLPVPWPTLVTTLTYGVQCFSRYPDHMKQHDFFKSAMPEGYVQERTIFFKDDGNYKTRA<br>EVKFEGDTLVNRIELKGIDFKEDGNILGHKLEYNYNSHNVYIMADKQKNGIKVNFKIRHNIED<br>GSVQLADHYQQNTPIGDGPVLLPDNHYLSTQSALSKDPNEKRDHMVLLEFVTAAGITLGM<br>DELYK |

Color denotation:

| | |
|---|---|
| (green) | eGFP |
| (orange) | Markers |
| (light purple) | Myc affinity tag |
| (grey) | HA affinity tag |

**SI References.**
